# Turnover of *PPP1R15A* mRNA encoding GADD34 controls the molecular memory of the cellular integrated stress response

**DOI:** 10.1101/2023.11.11.566686

**Authors:** Vera Magg, Alessandro Manetto, Katja Kopp, Chia Ching Wu, Mohsen Naghizadeh, Doris Lindner, Lucy Eke, Julia Welsch, Stefan M. Kallenberger, Johanna Schott, Volker Haucke, Nicolas Locker, Georg Stoecklin, Alessia Ruggieri

## Abstract

The integrated stress response (ISR) is a key cellular signaling pathway activated by environmental alterations that represses protein synthesis to restore homeostasis. To prevent sustained damage, the ISR is counteracted by the upregulation of Growth Arrest and DNA Damage inducible 34 (GADD34), a stress-induced regulatory subunit of protein phosphatase 1 that mediates translation reactivation and stress recovery. Here, we uncover a novel ISR regulatory mechanism that post-transcriptionally controls the stability of *PPP1R15A* mRNA encoding GADD34. We established that the 3’ untranslated region of *PPP1R15A* mRNA contains an active AU-rich element recognized by proteins of the ZFP36 family, promoting its rapid decay under normal conditions and stabilization for efficient expression of GADD34 in response to stress. We identify the tight temporal control of *PPP1R15A* mRNA turnover as a key step in establishing the ISR molecular memory, which sets the threshold for cellular responsiveness and mediates adaptation to repeated stress conditions.

## Introduction

The integrated stress response (ISR) is a highly sensitive signaling network that allows cells to quickly react and adapt to a variety of potentially harmful perturbations, including cellular damage, environmental stress and viral infections. Signals are transduced by four different kinases, namely heme-regulated inhibitor (HRI), general control nonderepressible 2 (GCN2), protein kinase R (PKR) and PKR-like endoplasmic reticulum kinase (PERK), whose activation results in the direct phosphorylation of a common downstream substrate, the alpha subunit of the eukaryotic translation initiation factor 2 (eIF2α)^1^. This step reduces the availability of the active Met-tRNA_i_^Met^/eIF2/GTP ternary complex and leads to stalling of translation initiation followed by ribosome run-off and polysome disassembly^2^. As a result, high local concentrations of released mRNAs and RNA-binding proteins trigger cytosolic phase separation and thereby drive the condensation of stress granules (SGs)^3^. In parallel, translational attenuation by increased levels of phosphorylated eIF2α (p-eIF2α) promotes the expression of *PPP1R15A* encoding Growth Arrest and DNA Damage inducible 34 (GADD34), a stress-induced adaptor protein that in complex with protein phosphatase 1 (PP1) mediates the dephosphorylation of eIF2α^4–6^. This negative feedback loop antagonizes prolonged activation of the ISR, allowing resumption of translation and return to homeostasis^7^.

Under normal conditions, maintenance of eIF2α in its non-phosphorylated form is essential for continuous translation. This is primarily achieved by the constitutive repressor of eIF2α phosphorylation (CReP), an alternative adaptor protein that targets PP1 to eIF2α^8^. In the absence of stress, *PPP1R15A* mRNA is expressed at low levels and is translationally repressed by two non-overlapping upstream open frames (uORFs) within the 5’ untranslated region (UTR). Under stress conditions, when the availability of active ternary complex is limiting, uORF translation ceases and ribosomal read-through allows translation of the *PPP1R15A* main ORF^9,10^. At the same time, two stress-responsive transcription factors, the activating transcription factor 4 (ATF4) and the C/EBP homologous protein (CHOP), are upregulated and induce transcription of the *PPP1R15A* gene^11–13^. Throughout this process, GADD34 is short-lived as a result of proteasomal degradation, enabling tight temporal control of its levels^14–16^.

We have previously described a unique example of a highly dynamic stress response to chronic infection with hepatitis C virus (HCV), characterized by alternating phases of SG assembly and disassembly, which coincide with phases of stalled and active translation, respectively^17^. By integrating quantitative experiments with mathematical modelling, we established that the molecular network generating this dynamic ISR is regulated by a stochastic process with memory, whereby cells switch to phases of SG assembly based on their previous experience. While we identified stochastic bursts of GADD34 expression and rapid GADD34 turnover as important determinants of this unique temporal control, we also found that *PPP1R15A* mRNA levels remain elevated for a prolonged period after stress release, beyond the time when transcription is turned off. *PPP1R15A* mRNA levels gradually return to their initial level over a period of approximately 20 hours, which delineates a window of cell adaptation to stress. Together, these results suggested that *PPP1R15A* mRNA acts as a molecular memory of previous ISR activation, facilitating subsequent bursts of GADD34 synthesis and hence rapid changes in eIF2α phosphorylation levels when cells are exposed to prolonged or repeated stress conditions^18^. These findings raise the question whether active control of *PPP1R15A* mRNA stability might serve as an additional, post-transcriptional layer at which the ISR and stress adaptation are regulated.

The balance between transcription and decay determines the abundance of every mRNA^19^. mRNA turnover is frequently regulated by *cis*-acting elements in the 3’ UTR, which can enhance or repress the stability of mRNAs^20^. Among them, AU-rich elements (AREs) represent a major class of mRNA destabilizing elements^21^ whose recognition by different RNA-binding proteins (RBPs) causes rapid mRNA degradation^22–25^, suppresses translation^26,27^ or enhances transcript stability^28^. Rapid degradation of ARE-containing mRNAs is initiated through deadenylation by the CCR4/NOT complex^29^, followed by decay from the 3’ end through the exosome^25,30^ or from the 5’ end by decapping and the exoribonuclease XRN-1^31,32^.

The human tristetraprolin (TTP, also known as ZFP36) family comprises three closely related RBPs: TTP, Butyrate Response Factor 1 (Brf1, also known as ZFP36L1) and Brf2 (ZFP36L2), all of which bind to the ARE of their target mRNAs through two tandem CCCH-type zinc finger domains (TZDs)^33^. The activity of TTP and Brf1 is regulated by inhibitory phosphorylation at several serine residues upon activation of the mitogen-activated protein kinase p38 (p38-MAPK)/MK2 pathway^34–37^. Brf1 is additionally phosphorylated at serine residue 92 by the phosphatidylinositol 3-kinase (PI3K)/Akt pathway^38,39^. Phosphorylation of TTP and Brf1 prevents ARE-mediated mRNA decay and enables efficient translation of target mRNAs^27,37–40^.

Here, we investigated the post-transcriptional regulation of *PPP1R15A* mRNA stability and uncovered its role in regulating the ISR. We identified a functional ARE in the *PPP1R15A* 3’ UTR, which is recognized by TTP and Brf1, and promotes rapid *PPP1R15A* mRNA decay in absence of stress. Exposure to different types of stress simultaneously activates the p38-MAPK/MK2 and PI3K/Akt pathways, leading to downstream inactivation of TTP and Brf1 decay activity, as well as to the rapid, transient stabilization of *PPP1R15A* mRNA. Our findings reveal that regulation of *PPP1R15A* mRNA stability controls the cellular sensitivity to stress and the kinetics of stress adaptation.

## Results

### Exposure to stress enhances *PPP1R15A* mRNA stability

We hypothesized that in addition to regulation at the transcriptional and translational level, *PPP1R15A* mRNA degradation rates may provide additional control of the dynamic response to HCV infection^18^. To examine this possibility, we first measured *PPP1R15A* mRNA turnover in Huh7 cells under normal conditions by actinomycin D chase experiments and quantitative real-time PCR (RT-qPCR), using the stable *GAPDH* mRNA as a reference. Intriguingly, we found *PPP1R15A* mRNA to be short-lived compared to *GAPDH* mRNA, with a half-life of approximately 1.2 hours (t_1/2_, Figure 1A). Upon chronic infection with HCV, *PPP1R15A* mRNA was weakly stabilized (t_1/2_ = 1.6 hours, Figure 1B left panel). This effect was stronger when the infection occurred in the presence IFN-α (t_1/2_ = 2.4 hours, Figure 1B right panel), consistent with our previous observation of a more dynamic SG response in infected cells treated with IFN-α^18^. We next investigated whether *PPP1R15A* mRNA stabilization might also occur in response to infection with dengue virus (DENV), a closely related RNA virus whose course of infection is acute, in contrast to HCV. Stabilization of *PPP1R15A* mRNA was observed early during infection (8 hpi t_/2_ = 1.4 hours, Figure 1C) and became more pronounced at a later stage (30 hpi t_1/2_ > 10 hours, Figure 1C), suggesting a dynamic regulation of *PPP1R15A* mRNA turnover over time. Despite different infection courses, the two viruses share a common feature in that they generate double-stranded (ds) RNA during genome replication. Transfection of cells with synthetic dsRNA also led to a strong stabilization (t_1/2_ > 10 hours, Figure 1D), indicating that the regulation of *PPP1R15A* mRNA turnover is triggered by the detection of dsRNA rather than by viral proteins.

**Figure 1.**
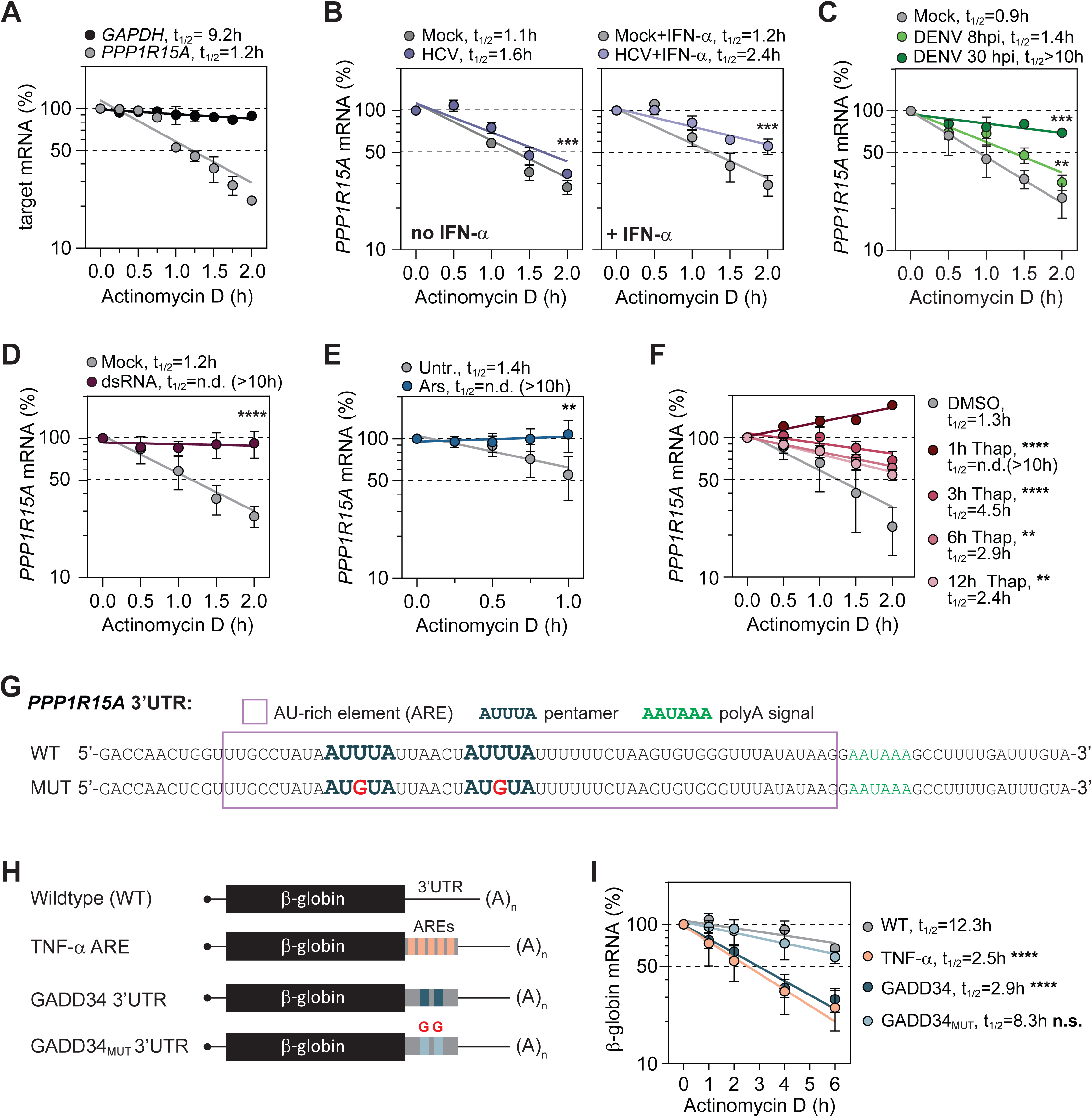
*PPP1R15A* 3’ UTR contains an RNA destabilizing AU-rich element (ARE). **(A)** Measurement of basal *PPP1R15A* mRNA decay. Levels of *PPP1R15A* and *GAPDH* mRNAs were determined following actinomycin D chase by RT-qPCR and normalized to a *firefly* luciferase spike-in RNA. *PPP1R15A* and *GAPDH* mRNA half-lives (t_1/2_) were calculated using a semi-logarithmic curve fitting. Shown are means ± SD of 3 biological replicates (n=3). **(B-F)** Decay measurements in Huh7 cells subjected to different types of stress (n=3): (B) infection with HCV for 48 h and subsequent treatment with IFN-α for 24 h, (C) infection with DENV for 8 and 30 h, (D) transfection with dsRNA for 16 h, (E) treatment with arsenite (Ars), and (F) treatment with thapsigargin (Thap) for up to 12 h. For all measurements, *PPP1R15A* mRNA levels were normalized to stable *GAPDH* mRNA levels. Statistical significance over all time points compared to mock, untreated (untr.) or DMSO controls is indicated. **(G)** Ribonucleotide sequence of the 3’ UTR of the human *PPP1R15A* mRNA. Wild type sequence is displayed on the top (WT). Sequence with mutated pentamers is displayed on the bottom (MUT). Purple box: ARE; AUUUA: ZFP36/TTP pentameric recognition motif; AAUAAA: polyA signal. **(H and I)** β-globin reporter mRNA turnover. Shown is a schematic of the reporters (H) in which the 3’ UTR of *PPP1R15A* (GADD34_WT_ or GADD34_MUT_) was inserted upstream of the β-globin 3’ UTR. TNF-α ARE served as a positive control. (I) β-globin reporter mRNA decay measurement in Huh7 stable cell lines (n=3). Statistical significance compared to WT β-globin reporter mRNA is indicated.

We next examined *PPP1R15A* mRNA turnover in response to the activation of different eIF2α-kinases by potent chemical stress inducers. We first chose sodium arsenite, which induces oxidative stress through the activation of HRI^41,42^. Remarkably, acute treatment of cells with arsenite led to a strong stabilization of *PPP1R15A* mRNA (t_1/2_ > 10 hours, Figure 1E). We then turned to PERK-mediated ER stress induced by thapsigargin treatment^43^, which cells can survive for more than 12 hours, to examine *PPP1R15A* mRNA turnover over a longer period of stress (Figure 1F). Thapsigargin treatment strongly stabilized *PPP1R15A* mRNA within the acute phase of the stress response, i.e. the first three hours of treatment (1h Thap, t_1/2_ > 10 hours, Figure 1F). Thereafter, as the stress response progressed toward the adaptation phase, *PPP1R15A* mRNA half-life gradually decreased over time (3h Thap, t_1/2_ = 4.5 hours, and 6h Thap, t_1/2_ = 2.9 hours), reaching at 12 hours (12h Thap, t_1/2_ = 2.4 hours) a value that remained significantly higher than that measured under normal conditions (DMSO, t_1/2_ = 1.3 hours) (Figure 1F). This finding supports our previous prediction of a progressive decrease in *PPP1R15A* mRNA levels during the stress adaptation phase, requiring more than 12 hours to return to basal levels^18^. Since *PPP1R15A* mRNA turnover is dynamically regulated in response to ISR activation by different triggers, *PPP1R15A* mRNA stabilization appears to be a conserved mechanism that contributes to the ISR negative feedback loop.

### *PPP1R15A* mRNA turnover involves a destabilizing AU-rich element in the 3’ untranslated region

Targeted control of mRNA degradation is frequently mediated by RBPs that recognize specific binding sites located in the 3’ UTR of mRNAs^44^. With 85 nucleotides, the 3’ UTR of human *PPP1R15A* mRNA is particularly short and comprises 70 percent adenosine (A) and uridine (U) residues (Figure 1G). Further inspection of the GADD34 3’ UTR sequence identified two canonical AUUUA pentamers embedded in U-rich clusters, which is typical for AREs as a major class of mRNA destabilizing elements^21,45^. To examine whether *PPP1R15A* mRNA turnover is regulated via a functional ARE, we generated stable Huh7 cell lines expressing a rabbit β-globin reporter gene^46^ in which we inserted, between the stop codon and the β-globin 3’ UTR, either the wild type *PPP1R15A* 3’ UTR sequence (GADD34) or a mutated sequence where both AUUUA pentamers were converted to AUGUA (GADD34_MUT_) (Figure 1H). As positive control, we used a β-globin reporter gene harboring the ARE of tumor necrosis factor-α (TNF-α) known to strongly accelerate mRNA decay^22,37,46^. Remarkably, insertion of the *PPP1R15A* 3’ UTR promoted the decay of the otherwise stable β-globin reporter mRNA (WT, t_1/2_ = 12.3 hours; GADD34, t_1/2_ = 2.9 hours) almost as efficiently as the TNF-α ARE (t_1/2_ = 2.5 hours) (Figure 1I). Mutation of the two AUUUA pentamers prevented rapid mRNA turnover (GADD34_MUT_, t_1/2_ = 8.3 hours), confirming the presence of an active ARE with destabilizing activity in the *PPP1R15A* 3’ UTR.

### *PPP1R15A* ARE regulates mRNA engagement with polysomes

To consolidate this result, we sought to explore the impact of deleting the ARE from the endogenous *PPP1R15A* locus, within the third exon of the *PPP1R15A* gene (Figure S1A). We generated Huh7 GADD34ΔARE single-cell clones using the CRISPR-Cas9 technology and nucleofection of two CRISPR RNAs (crRNAs) targeting the regions immediately upstream and downstream of the ARE. Three homozygous KO cell clones were selected by analysis of the *PPP1R15A* genomic sequence (Figure S1A). Additionally, control (Ctrl) cell clones were generated by nucleofection of non-targeting crRNAs. We analyzed *PPP1R15A* mRNA levels at steady state and upon treatment with thapsigargin. While deletion of the ARE did not significantly alter basal mRNA levels (Figure 2A), it resulted in a stronger increase of *PPP1R15A* mRNA levels in response to thapsigargin exposure during the stress adaptation phase, particularly at 6 hours post treatment (Figure 2B). This result was supported by the slower decay of *PPP1R15A* mRNA measured during stress adaptation, after treatment with thapsigargin for 3 hours (Figure 2C, 3h Thap: Ctrl t_1/2_ = 1.7 hours, ΔARE t_1/2_ >10 hours). Interestingly, Huh7 GADD34ΔARE cells showed a modest though significant increase in the basal GADD34 mRNA half-life (Figure 2C, DMSO: Ctrl t_1/2_ = 0.9 hours, ΔARE t_1/2_ = 1.2 hours).

**Figure 2.**
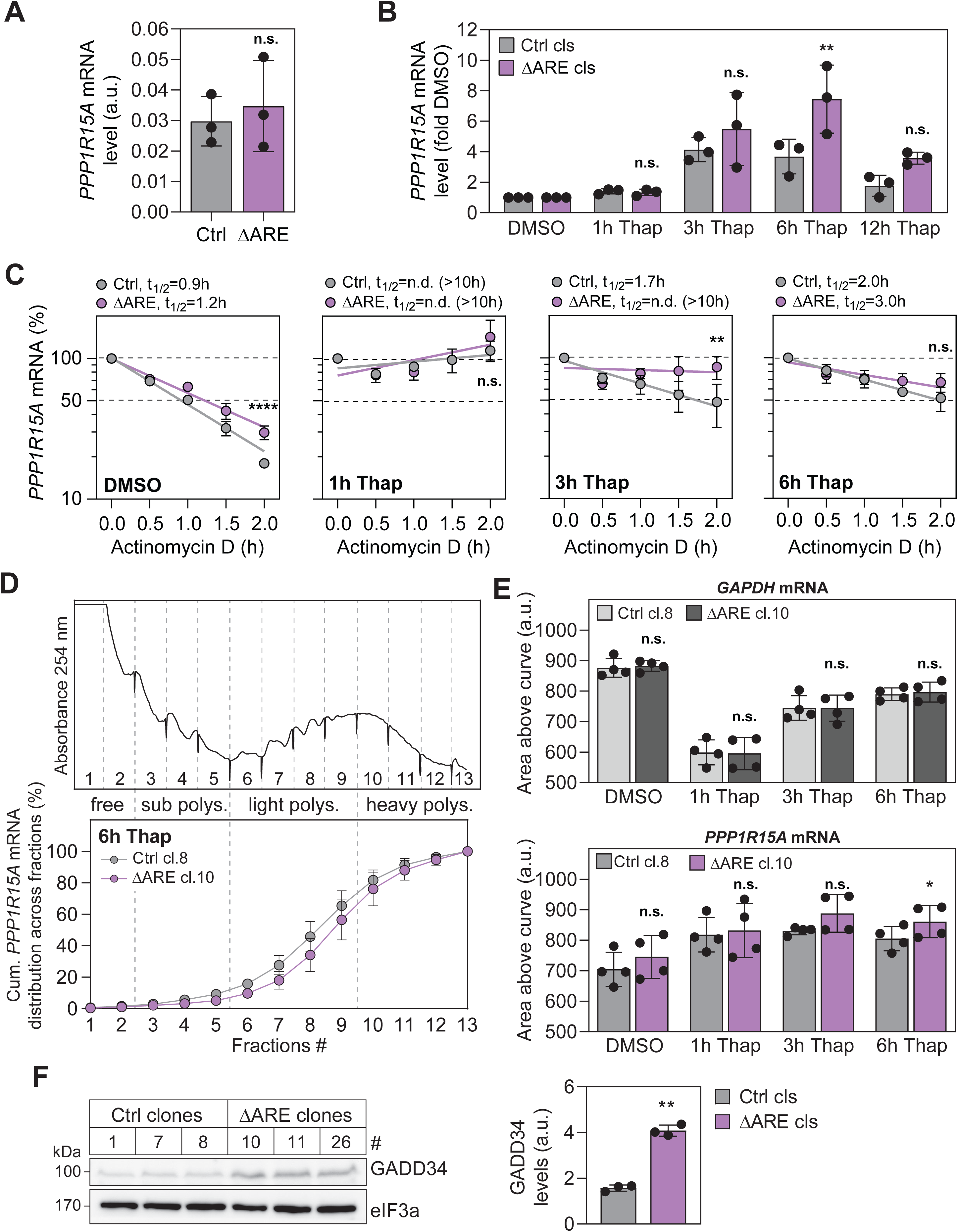
*PPP1R15A* ARE regulates mRNA engagement with polysomes. **(A)** Measurement of relative basal *PPP1R15A* mRNA expression levels in Huh7 control (Ctrl) and GADD34ΔARE (ΔARE) cell clones. Values were normalized to stable *GAPDH* mRNA levels. Shown is the representative mean value ± SD of three Ctrl and three ΔARE individual cell clones (n=2). **(B)** *PPP1R15A* mRNA expression levels in Huh7 Ctrl and ΔARE clones (cls) after treatment with DMSO or thapsigargin for up to 12 h. Values were normalized to GAPDH mRNA levels and are depicted as fold DMSO controls. Histograms show the mean ± SD of three Ctrl and three ΔARE individual cell clones (cls) (n=1). Statistical significance compared to Ctrl cells at the individual time points is indicated. **(C)** *PPP1R15A* mRNA decay in Huh7 Ctrl and ΔARE cell clones (cls). Cells were treated with thapsigargin or DMSO control for up to 6 h prior to addition of actinomycin D. Shown is the mean ± SD of three Ctrl and three ΔARE individual cell clones (n=1). Statistical significance over all time points compared to Ctrl is indicated. **(D and E)** Polysome profiles of Huh7 Ctrl cl.8 and Huh7 GADD34ΔARE cl.10 treated with thapsigargin for up to 6 h were recorded followed by fractionation of the sucrose density gradients. Shown are mean ± SD of two technical replicates from two distinct biological replicates (n=2). Total RNA was extracted from the polysome fractions, and the distribution of *PPP1R15A* and *GAPDH* mRNA across the polysome gradients was analyzed by RT-qPCR. (D) Shown is a representative profile (top panel). The mean percentage ± SD of mRNA in each fraction is depicted as a cumulative distribution. A representative cumulative curves for *PPP1R15A* mRNA is shown in the bottom panel. (E) For each condition, the area above the curve was calculated as a measure for the overall association of the mRNA with ribosomes. Statistical significance compared to Ctrl clones is indicated. **(F)** Western blot analysis of GADD34 basal expression in three Huh7 Ctrl and three GADD34ΔARE individual cell clones (n=1) (left panel). Quantification is shown on the right panel. Measured GADD34 band intensities were normalized to eIF3a and are depicted as arbitrary units (a.u.). Statistical significance compared to Ctrl clones is indicated.

Since AREs are also known to attenuate translation^26,27^, we tested whether ARE deletion affects the *PPP1R15A* mRNA translation efficiency, and performed polysome profile analyses with Huh7 Ctrl and GADD34ΔARE cells treated with thapsigargin for different periods of time (Figures 2D, S1B and S1C). The distribution of *GAPDH* and *PPP1R15A* mRNAs across the polysome profile fractions was analyzed by RT-qPCR (Figures S1B and S1C). As anticipated for the mRNA of a housekeeping gene, *GAPDH* mRNA (Figure S1B) was enriched in the heavy polysomal fractions under basal conditions (DMSO). During the acute phase of stress (1h Thap), it shifted towards the monosomal and light polysomal fractions, reflecting substantial translational repression. Entering the stress adaptation phase (3 and 6h Thap), *GAPDH* mRNA progressively re-engaged with heavy polysomal fractions as global translation resumed. Importantly, *GAPDH* mRNA distribution in polysomes (Figure S1B) as well as its proportion associated with polysomes - estimated by calculating the area above the cumulative mRNA distribution (Figure 2E, top panel) - was identical in Ctrl and GADD34ΔARE cells. This result indicated that levels of global translation were similar in both cell lines.

In contrast to *GAPDH*, *PPP1R15A* mRNA showed the opposite pattern typical for a stress-induced protein (Figure S1C). Under basal conditions (DMSO), *PPP1R15A* mRNA was mainly associated with light polysomal fractions, in line with its translation being attenuated by two uORFs^10^. Upon deletion of the ARE, the mRNA exhibited a slight but highly reproducible shift towards the heavy polysomal fractions (DMSO, Figures 2E bottom panel and S1C). While the difference in the proportion of mRNA associated with polysomes was not significant (DMSO, Figure 2E bottom panel), the observed shift in polysome distribution was consistent with the fact that ARE deletion led to a stronger increase in GADD34 levels (Figure 2F) than in *PPP1R15A* mRNA levels (Figure 2A).

Upon treatment with thapsigargin, both *PPP1R15A* and *PPP1R15AΔARE* mRNA shifted towards heavier polysomal fractions, reflecting their translational de-repression. During stress adaptation, *PPP1R15A* mRNA gradually shifted back to the lighter polysomal fractions (6h Thap, Figures 2E and S1C), indicating a slow return to the initial state of repression. Notwithstanding, *PPP1R15AΔARE* mRNA remained more engaged with heavier polysomal fractions than *PPP1R15A* mRNA during stress adaptation (6h Thap, Figures 2E and S1C). Altogether, these results suggest that the engagement of *PPP1R15A* mRNA with polysomes correlated with its reduced turnover in Huh7 GADD34ΔARE cell clones (Figure 2C), indicating that by acting on the turnover and translation of *PPP1R15A* mRNA, the ARE is important for controlling GADD34 levels under basal conditions and during stress adaptation.

### The pre-existing pool of *PPP1R15A mRNA* is stabilized during the acute phase of the ISR

The precise temporal control of signaling pathways is crucial for rapid responses to changing environmental cues, impacting cell fate decisions. Expressing a pool of mRNA that is kept translationally repressed in the absence of stress may enable cells to respond more promptly to changes in eIF2α phosphorylation levels before transcriptional induction. To determine whether the pre-existing and/or newly synthesized *PPP1R15A* mRNA pools are stabilized during the acute phase of the stress response, we employed two complementary approaches (Figure 3A).

**Figure 3.**
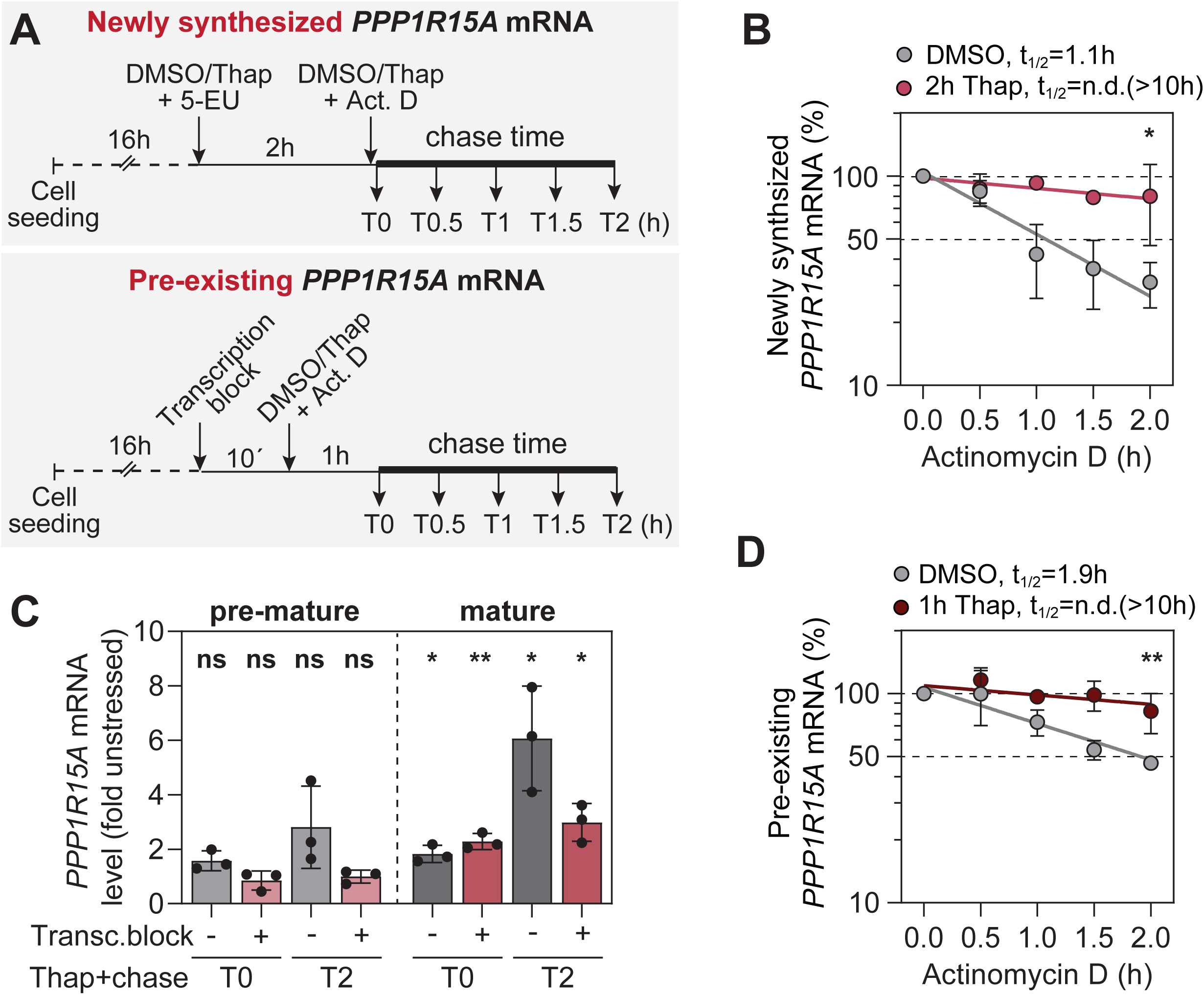
The pre-existing pool of *PPP1R15A mRNA* is stabilized during the acute phase of the ISR. **(A)** Experimental design. For the analysis of the turnover of *PPP1R15A* newly synthesized mRNA (top panel), Huh7 cells were co-treated with 5-EU and DMSO or thapsigargin (Thap) for 2 h prior to a 2-hour actinomycin D chase. Cells were harvested at the indicated time points and 5-EU-labelled mRNA biotinylated by click-it chemistry and further purified on beads. For the analysis of the turnover of *PPP1R15A* pre-existing mRNA (bottom panel), Huh7 cells were treated with actinomycin D for 10 min to block transcription prior to a 1-hour treatment with DMSO or thapsigargin, and followed by a 2-hour actinomycin D chase. Cells were harvested at the indicated time points. **(B)** Decay measurement of the newly synthesized *PPP1R15A* mRNA (n=3). Levels of *PPP1R15A* mRNA were normalized to T0 and plotted as percentage of T0. Statistical significance over all time points compared to DMSO is indicated. **(C and D)** Analysis of pre-existing *PPP1R15A* mRNA turnover in Huh7 cells treated with thapsigargin for 1 h (n=3). (C) Measurement of *PPP1R15A* pre-mRNA and mature mRNA levels in Huh7 cells treated with DMSO or thapsigargin, at the start of the actinomycin D chase experiment (T0) and at the end (T2). Levels of *PPP1R15A* mRNA were normalized to GAPDH mRNA and plotted as percentage of T0. Values are shown as fold DMSO. Shown is the mean ± SD (n=3). Statistical significance compared to DMSO-treated cells is indicated. (D) Decay measurement of the pre-existing *PPP1R15A* mRNA in Huh7 cells treated with thapsigargin (n=3). Shown is the mean ± SD and statistical significance compared to DMSO.

First, the newly synthesized *PPP1R15A* mRNA (Figure 3A, top panel) was metabolically labelled with 5-ethynyluridine (5-EU), an alkyne-modified uridine that can subsequently be linked to biotin by click-chemistry, in a 2-hour co-treatment of with thapsigargin or DMSO, prior to the start of the actinomycin D chase. At each time point, total RNA was extracted, 5-EU was clicked to biotin, and newly synthesized RNA purified using streptavidin beads. The amount of *PPP1R15A* mRNA bound to the beads was measured by RT-qPCR. Similar to the results obtained with the overall pool of *PPP1R15A* mRNA (Figure 1F), the newly synthesized *PPP1R15A* mRNA was short-lived, with a half-life of approx. 1.1 hours in DMSO-treated cells (Figure 3B), and strongly stabilized upon thapsigargin treatment (Thap 2h, t_1/2_ > 10 hours).

In a second approach, we addressed the fate of the pre-existing pool of *PPP1R15A* mRNA (Figure 3A, bottom panel. To this end, cells were treated with actinomycin D prior to stress exposure to prevent *de novo* transcription of *PPP1R15A* mRNA in response to ISR activation (Figure 3C). Cells were then subjected to 1-hour thapsigargin treatment followed by actinomycin D chase. In the absence of stress, the pre-existing *PPP1R15A* mRNA had a rapid turnover (t_1/2_ = 1.9 hours), while it was strongly stabilized in the acute phase of the stress response to thapsigargin (t_1/2_ >10 hours, Figure 3D). This result indicates that stabilization of the pre-existing *PPP1R15A* mRNA pool during the acute phase of the ISR represents the first regulatory step promoting rapid GADD34 production, before its transcriptional and translational induction.

### *PPP1R15A* ARE is recognized by RNA-binding proteins of the ZFP36 family

Next, we interrogated the *PPP1R15A* 3’ UTR for destabilizing RBP binding sites using RBPDB, a database that integrates sequence information from *in vitro* and *in vivo* RNA-binding experiments^47^. The nonameric sequence CUAUUUAUU (Figure 1G) was highly scored as a potential binding site for TTP (ZFP36), a prediction confirmed by the AREsite2 database^48^. ZFP36 family proteins represent the most potent inducers of ARE-mRNA decay^49^, and comprise three members, TTP, Brf1 and Brf2, all of which contain a CCCH-type TZD with high affinity for AREs^50^. The TZF domains of ZFP36 proteins share 70 percent sequence identity, and there is evidence that TTP and Brf1 family members share some common target mRNAs ^51^ but also have preferences for specific ARE-mRNAs^52,53^. We therefore investigated both TTP and Brf1. In line with its function as an immediate early gene^54,55^, the basal expression of *ZFP36* mRNA was low at steady state compared to *ZFP36L1* mRNA (Figure S2A), but transiently induced by exposure to thapsigargin (Figure S2B).

First, we assessed the ability of the two ZFP36 family members to bind to endogenous *PPP1R15A* mRNA under normal conditions. To overcome the difficulty of detecting endogenous TTP and Brf1 in Huh7 cells (Figure S2C), we transiently expressed myc-his-tagged human TTP and Brf1 (Figure 4A) at low levels to avoid the spontaneous formation of SGs^37,56^. We also included the TTP murine orthologue (mTTP) and its previously characterized RNA-binding mutant, harboring a CCCH to SSCH mutations in both TZF domains (mTTP_m1,2_)^29^. Myc-his-tagged GFP served as control for unspecific RNA binding. In contrast to *GAPDH* mRNA, endogenous *PPP1R15A* mRNA was efficiently immunoprecipitated with TTP, mTTP and Brf1, but not mTTP_m1,2_ (Figure 4B), confirming the importance of the TZF domain in the interaction of ZFP36 proteins with *PPP1R15A* mRNA.

**Figure 4.**
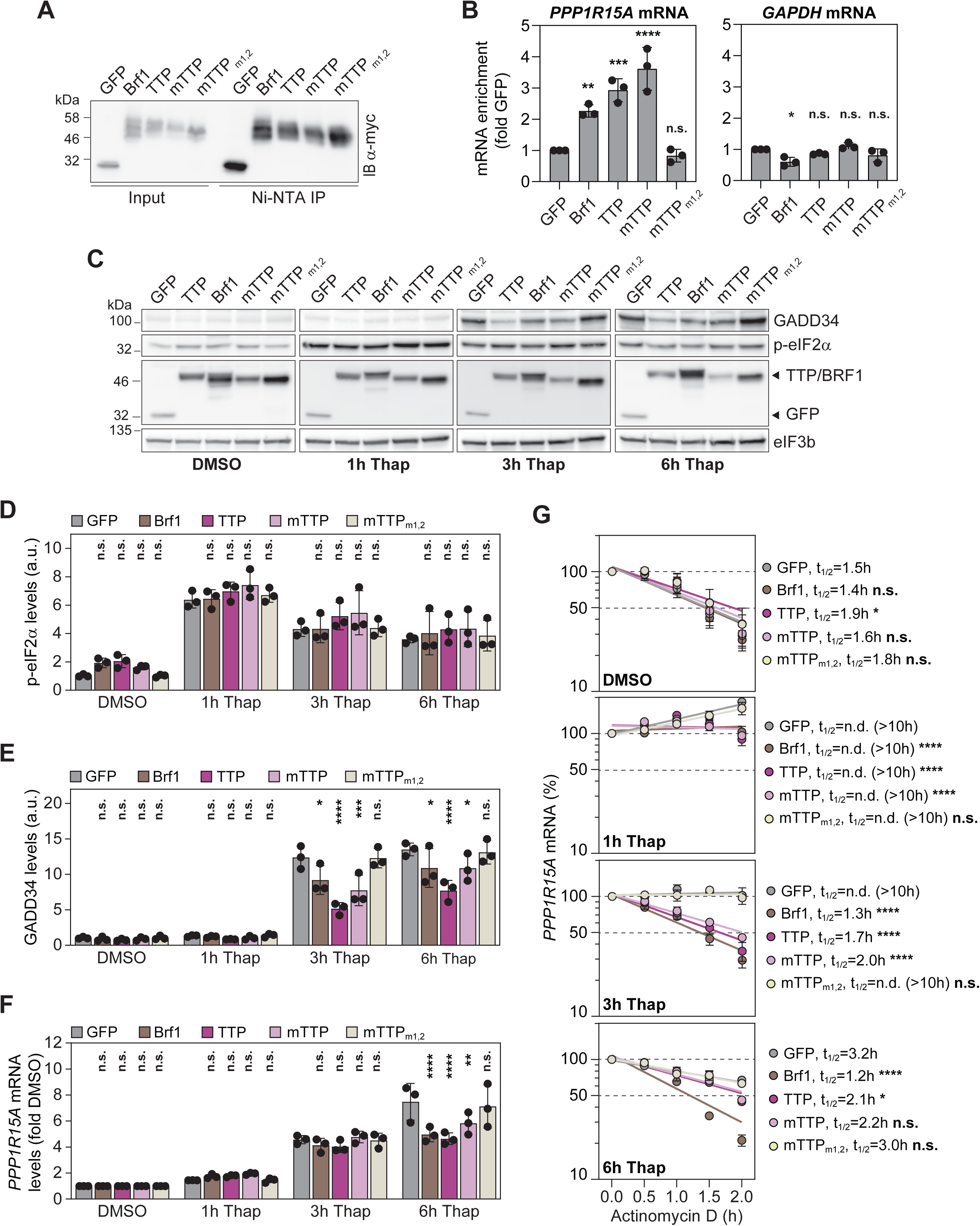
Overexpression of TTP and Brf1 induces *PPP1R15A* mRNA destabilization during stress adaptation. **(A)** Myc-his-tagged GFP, TTP, Brf1, TTP murine orthologue (mTTP) or RNA-binding mutant (mTTP_m1,2_) were immunoprecipitated using Ni-NTA beads (Ni-NTA IP) from transiently expressing Huh7 cells. RNA was subsequently isolated and analyzed by RT-qPCR (n=3). **(B)** Analysis of co-precipitated *PPP1R15A* and GAPDH mRNAs, normalized to GFP control. Statistical significance compared to GFP is indicated. **(C-G)** Transient overexpression of myc-his tagged proteins in Huh7 cells treated with DMSO or thapsigargin for the indicated times. (C) Shown is a representative Western blot analysis (n=3). Corresponding Western blot quantifications (mean ± SD) of p-eIF2α (D) and GADD34 (E) levels. Measured band intensity values were normalized to eIF3b and are depicted as arbitrary units (a.u.) (n=3). Statistical significance compared to GFP at each time point is indicated. (F) Corresponding quantification of *PPP1R15A* mRNA levels by RT-qPCR. *PPP1R15A* mRNA levels were normalized to *GAPDH* mRNA values and are represented as fold of DMSO (n=3). Statistical significance compared to GFP at each time point is indicated. (G) *PPP1R15A* mRNA decay in cells treated with DMSO or thapsigargin. Levels of *PPP1R15A* mRNA were normalized to stable GAPDH mRNA and plotted as percentage of T0. Shown is the mean ± SD (n=3). Statistical significance over all time points compared to GFP is indicated.

Although predominantly located in 3’ UTRs, AREs are also distributed in the intronic regions of some pre-mRNAs^57^. To ensure that ZFP36 proteins specifically bind to the ARE located in the 3’ UTR of *PPP1R15A* mRNA, we performed similar experiments using the β-globin reporter cell lines transiently expressing myc-his-tagged RBPs (Figure S2D). The reporter mRNA harboring the *PPP1R15A* 3’UTR was strongly enriched following immunoprecipitation of TTP, Brf1 and mTTP, but not mTTP_m1,2_ (Figure S2E). Despite being significantly reduced, the binding of the reporter mRNA harboring the GADD34_MUT_ 3’UTR was not completely abrogated, suggesting that other sequences or secondary structures outside of the two AUUUA pentamers may contribute to the binding of ZFP36 proteins to GADD34 mRNA. Together, these results identified the ZFP36 members TTP and Brf1 as *PPP1R15A* ARE-interacting proteins.

### Ectopic expression of TTP and Brf1 destabilizes *PPP1R15A* mRNA

Previous studies showed that TTP and Brf1 recruit the mRNA decay machinery to selectively degrade ARE-mRNAs^29,58^. We therefore investigated the impact of ectopically expressed TTP and Brf1 on *PPP1R15A* mRNA turnover and GADD34 expression under stress conditions using thapsigargin for different periods of time (Figure 4C). Interestingly, TTP and Brf1 transient overexpression appeared to slightly, but not significantly, increase basal levels of eIF2α phosphorylation (p-eIF2α) in the absence of stress (DMSO, Figure 4D). Levels of p-eIF2α increased in the acute phase of stress in all cell lines (1h Thap, Figure 4D) and decreased during the stress adaptation phase (3h and 6h Thap, Figure 4D), mirroring the appearance of GADD34 (3h and 6h Thap, Figure 4E). Notably, GADD34 levels were markedly lower in cells expressing TTP, Brf1 or mTTP compared to those expressing GFP or mTTP_m1,2_ (Figures 4E). This reduction was only detectable at the mRNA level at 6h post thapsigargin treatment (Figure 4F), possibly due to the detection of partially deadenylated and/or degraded transcripts by RT-qPCR. Although TTP and Brf1 overexpression reduced GADD34 levels, it did not result in an apparent higher level of p-eIF2α. We therefore used SG formation as a more sensitive readout for subtle changes in p-eIF2α and confirmed that SG disassembly was delayed approximately 2.7-fold during the stress adaptation phase (3 hours post thapsigargin removal) in TTP and Brf1-overexpressing cells (Figures S3A and S3B).

To substantiate our results, we analyzed *PPP1R15A* mRNA decay over time in the different overexpressing cell lines in the absence (DMSO) and presence of thapsigargin. In control cells expressing GFP or msTTP_m1,2_, the decay kinetics of *PPP1R15A* mRNA was as previously observed in parental cells (Figure 1F), with a strong stabilization during the acutef phase of stress (1h Thap), followed by a gradual return of decay during the adaptation phase (Figure 4G, see also same data arranged differently in Figure S3C). This dynamic was altered in cells expressing TTP, Brf1 or mTTP, in which *PPP1R15A* mRNA returned faster to rapid decay during the stress adaptation phase (Figure 4G). Additionally, we used a constitutively active, non-phosphorylable S52A/S178A mutant of murine TTP, mTTP-AA, unable to bind to 14-3-3 isoforms^34,37^. *PPP1R15A* mRNA remained labile in presence of the TTP-AA mutant (Figures S4A and S4B), confirming that phosphorylation-dependent inhibition of ZFP36 proteins contributes to the stabilization of *PPP1R15A* mRNA during the acute phase of the stress response. While initially inactivated during the acute phase of the stress response, ZFP36 proteins accelerate the clearance of *PPP1R15A* mRNA during the adaptation phase of the stress response.

### Acute and chronic stresses activate the p38-MAPK/MK2 and PI3K/Akt signaling pathways

TTP and Brf1 activity is downregulated by their phosphorylation via the p38-MAPK/MK2 and PI3K/Akt signaling pathways^37,38^. We thus explored whether the stress response induced by acute treatment with thapsigargin or chronic HCV infection could represent a signal for MAPK activation using a commercially available phospho-MAPK array (Figure S5A). While this array detected a strong increase in Akt phosphorylation (p-Akt) in response to both acute thapsigargin treatment and chronic HCV infection, increased levels of p38α phosphorylation (p-p38α) were only detected upon thapsigargin treatment. However, using another set of antibodies and Western blot analyses, we confirmed that both pathways are activated by chronic HCV infection (Figures 5A and 5B) and treatment with thapsigargin (Figures S5B and S5C). Consistent with this observation, our previous work had shown that acute dengue virus infection induced the activation of both p38-MAPK/MK2 and PI3K/Akt pathways and increased GADD34 levels^59^.

**Figure 5.**
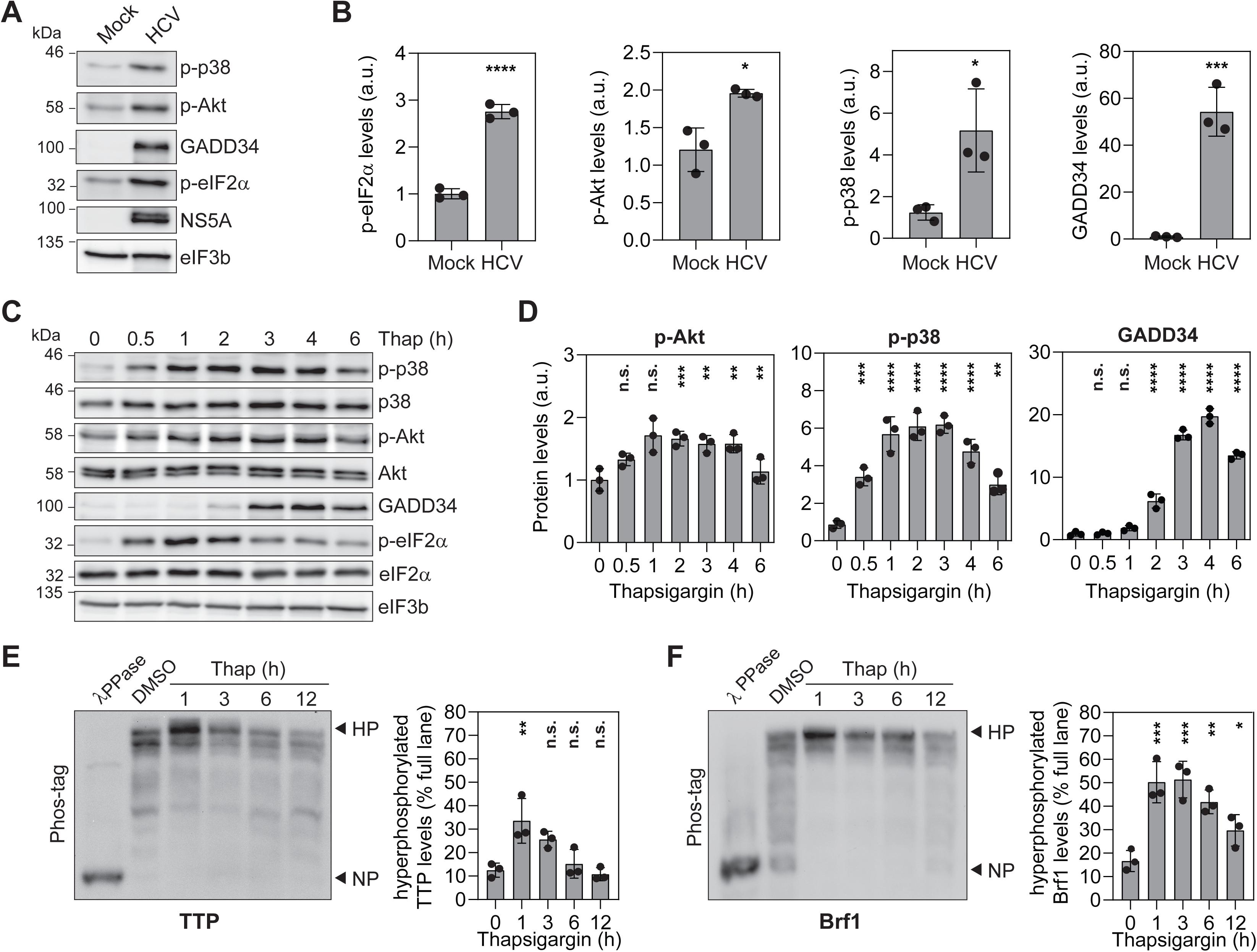
Stress-mediated activation of the p38-MAPK/MK2 and PI3K/Akt signaling pathways mediates TTP and Brf1 phosphorylation. **(A and B)** Analysis of p38-MAPK/MK2 and PI3K/Akt kinase signaling pathways upon HCV infection. Huh7 cells were infected with HCV for 48 h prior to the addition of IFN-α. At 72 h post infection, cell were harvested and lysates analyzed (n=3). (A) Shown is a representative Western blot. (B) Corresponding quantifications (mean ± SD) of p-eIF2α, p-p38, p-Akt, GADD34, and HCV NS5A levels. Measured band intensities were normalized to eIF3b and depicted as arbitrary units (a.u.). Statistical significance compared to Mock is indicated. **(C and D)** Dynamics of p38-MAPK/MK2 and PI3K/Akt pathway activation upon thapsigargin treatment. Huh7 cells were treated with DMSO or thapsigargin for up to 6 h (n=3). (C) Shown is a representative Western blot. (D) Corresponding quantifications (mean ± SD). Statistical significance compared to DMSO is indicated. **(E and F)** Analysis of TTP and Brf1 phosphorylation levels by Phos-tag acrylamide gel and Western blot (n=3). Cells transiently expressing myc-his-tagged TTP (E) or Brf1 (F) were treated with DMSO or thapsigargin for the indicated times. Cell extracts digested with lambda phosphatase (λPPase) were used to determine the migration pattern of non-phosphorylated forms. HP, hyper-phosphorylated form. NP, non-phosphorylated form. Shown are representative Western blot analyses (left panels) and corresponding quantifications (right panels). Signal was detected using a myc-tag antibody. Band intensities of the hyper-phosphorylated forms are depicted as percentage of the total signal intensity in the corresponding lane (mean ± SD). Statistical significance compared to DMSO (0) is indicated.

To obtain insight into the dynamics of p38-MAPK/MK2 and PI3K/Akt pathway activation, we analyzed the levels of p-p38 and p-Akt at different time points after treatment with thapsigargin (Figure 5C and 5D). Levels increased during the acute phase of the stress response within the first hour of treatment, preceding the induction of GADD34 expression, detectable 2 to 3 hours after treatment. During the stress adaptation phase, i.e. from 3 hours onwards, p-p38 and p-Akt levels started to decrease, concomitantly with the increase in GADD34 levels (Figures 5C and 5D). A similar dynamics of pathway activation was also observed with arsenite, within a shorter window of treatment that allowed cells to survive (Figures S5D and S5E). Taken together, these results show that the p38-MAPK/MK2 and PI3K/Akt signaling pathways are transiently activated during the ISR.

### Activation of the ISR leads to transient hyper-phosphorylation of ZFP36 proteins

Both TTP and Brf1 are multiply phosphorylated in cells^35,60,61^. We therefore assessed changes in the global phosphorylation levels of TTP and Brf1 over time in thapsigargin-treated cells. Extracts of myc-his tagged TTP and Brf1 expressing cells were separated by Phos-tag polyacrylamide gel electrophoresis (Figures 5E and 5F). The phosphorylation pattern was compared to that of the non-phosphorylated forms (NP), obtained by treatment of the extracts with lambda protein phosphatase (λPPase). As previously reported, the presence of multiple slower-migrating bands indicated that TTP and Brf1 are extensively phosphorylated in the absence of stress (0h). Stimulation with thapsigargin resulted in a noticeable shift of TTP and Brf1 towards a hyper-phosphorylated form (HP) in the acute phase of the stress response (1h). HP levels decreased continuously thereafter, as cells progressed to the stress adaptation phase, reaching almost basal levels 12 hours after treatment. The observed dynamic regulation of ZFP36 protein phosphorylation correlated with the activation kinetics of the PI3K/Akt and p38-MAPK/MK2 signaling pathways and coincided with the decreased decay activity measured during the acute phase of the stress response (1h Thap, Figure 4G). During the stress adaptation phase, dephosphorylation of ZFP36 proteins went along with the decreasing MAPK activity and decreasing GADD34 expression levels (6h Thap, Figures 5C and 5D). Altogether, these results suggest that ZFP36 proteins are induced yet their decay activity is transiently inactivated by hyperphosphorylation during the acute phase of the ISR, thereby enabling efficient GADD34 expression. During progression to the stress adaptation phase, gradual dephosphorylation of TTP and Brf1 promotes the timely degradation of *PPP1R15A* mRNA.

### ZFP36 proteins attenuate GADD34 expression in the absence of stress and during stress adaptation

To further evaluate the role of TTP and Brf1 in the post-transcriptional regulation of GADD34, we generated TTP and Brf1 KO cell clones using a CRISPR/Cas9 two-guide strategy, with crRNAs targeting the regions immediately upstream and downstream of the *ZFP36 and ZFP36L1* TZDs (Figure S6A and S6B). The deletion of the genomic DNA sequences between the two targeted sites was confirmed in the independent cell clones by PCR and sequencing. Control cell clones (Ctrl) were generated for comparison using a non-targeting crRNA. While *PPP1R15A* steady state mRNA levels appeared slightly elevated in the TTP and Brf1 KO cell clones, the differences were not statistically significant (Figure S6C). Similarly, there were only marginal effects on *PPP1R15A* mRNA turnover in the single TTP and Brf1 KO cell clones (Figures S6D and S6E). Likewise, GADD34 protein levels were barely affected by single TTP and Brf1 KO, and only TTP KO cells showed a 1.3-fold increase at 6 hours post thapsigargin treatment (Figure S6F).

Since the effects in the single KO clones were marginal, we explored the possibility of a functional redundancy by creating double KO (DKO) cell clones, in which TTP was knocked out in a Brf1 KO background (Figure S7A). Importantly, the analysis of transcriptome-wide changes in mRNA abundance between Huh7 Ctrl and DKO cells indicated that the depletion of both TTP and Brf1 did not result in a global defect in mRNA homeostasis, confirming that ZFP36 proteins act specifically on a very restricted group of target mRNAs, including *PPP1R15A* mRNA whose abundance was increased in DKO cells compared to Ctrl cells (Figure S7B). In contrast to the single KO clones, *PPP1R15A* mRNA was clearly more stable in the Brf1/TTP DKO clones (Figure 6A). This effect was reflected at the protein level, with a 1.8-fold increase in the basal expression of GADD34 in the DKO cell clones (Figure 6B) while global protein synthesis was not altered in these cells (Figure S7C).

**Figure 6.**
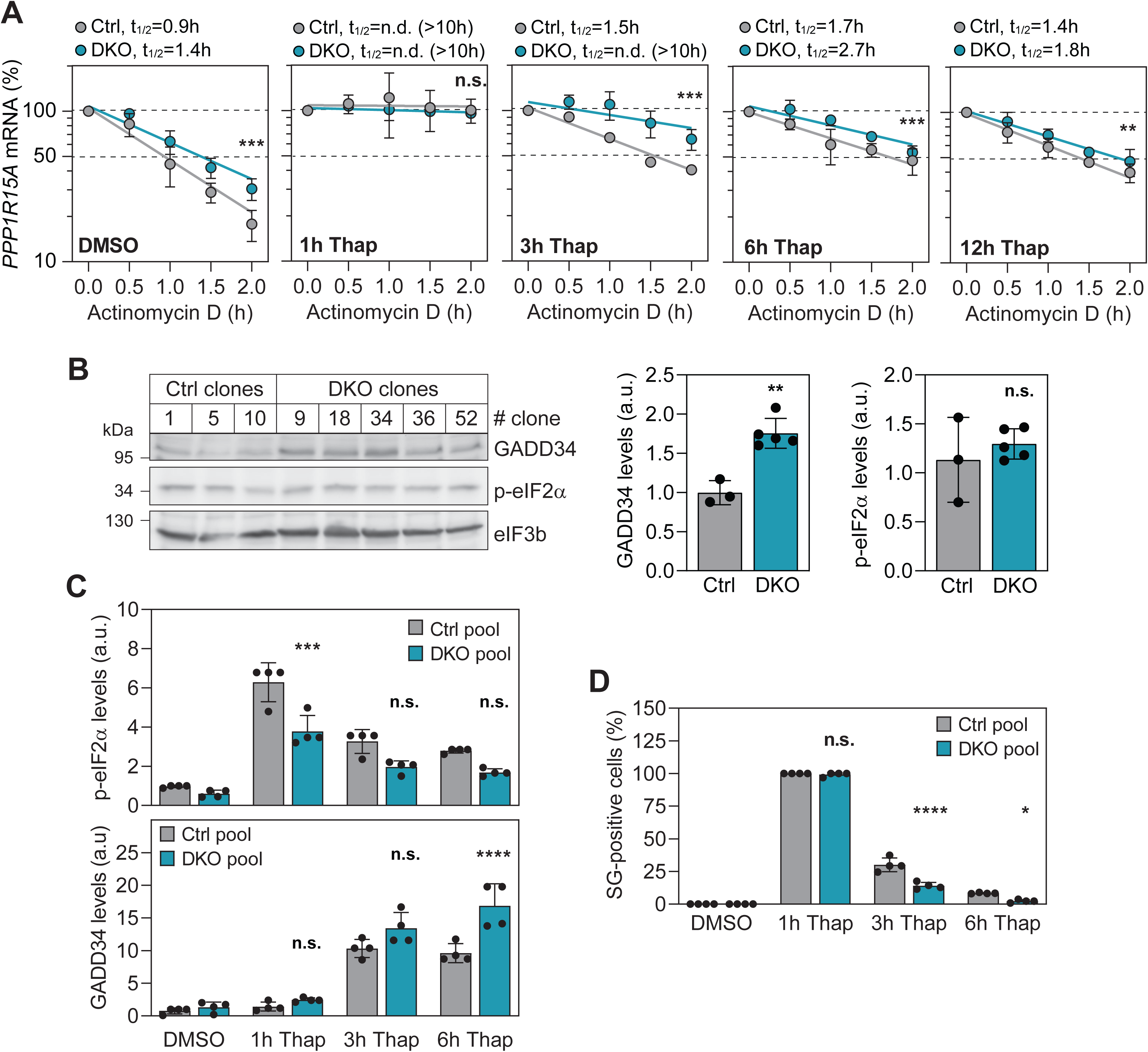
Increased basal GADD34 expression in Huh7 Brf1/TTP DKO cells. **(A)** *PPP1R15A* mRNA decay measurements in Huh7 control (Ctrl) and Brf1/TTP double knockout (DKO) cells. Cell pools were generated for Ctrl with cell clones 1, 5 and 10 and for DKO with cell clones 9, 18, 34, 36 and 52. Cell pools were treated with DMSO or thapsigargin for up to 12 h prior to 2-hour actinomycin D chase. Shown is the mean ± SD (n=3). Statistical significance over all time points compared to pooled Ctrl is indicated. **(B)** Analysis of GADD34 and p-eIF2α basal expression levels in Huh7 Ctrl and DKO individual cell clones. Shown is a representative Western blot analysis (top panel) and corresponding quantifications of GADD34 (bottom left panel) and p-eIF2α (bottom right panel) levels (n=2). **(C and D)** Analysis of GADD34 and p-eIF2α expression levels in Huh7 Ctrl and DKO cell pools treated with DMSO or with thapsigargin up to 6 h (C) and corresponding SG analysis (D). Measured band intensities were normalized to eIF3a and depicted as arbitrary units (a.u.). Shown is the mean ± SD (n=4). Statistical significance compared to Ctrl cell pools is indicated.

Enhanced stability of *PPP1R15A* mRNA was also observed in DKO cells upon thapsigargin treatment, especially during the early phase of stress adaptation (3h Thap, Figure 6A). This difference was also associated with increased GADD34 expression during the adaptation phase (6h Thap, Figures 6C, S7D and S7E), reduced phosphorylation of eIF2α (Figure 6C) and slower clearance of SGs (Figure 6D).

In light of these results, we hypothesize that TTP and Brf1 have redundant roles in regulating *PPP1R15A* mRNA turnover, maintaining low levels under normal conditions. In response to stress, when ZFP proteins regain their decay-promoting activity during the adaptation phase, they accelerate *PPP1R15A* mRNA clearance. Under stress conditions, TTP appears to be the dominant regulator of *PPP1R15A* mRNA stability, consistent with its more dynamic expression profile (Figure S2B).

### ZFP36 proteins contribute to setting the threshold for stress responsiveness and to the dynamics of stress adaptation

We previously found that the sustained presence of *PPP1R15A* mRNA contributes to the transient adaptation to stress, reducing the responsiveness to a second stress insult. Directly after release from acute stress, cells enter a refractory phase during which the remaining GADD34 protein dephosphorylates eIF2α, effectively repressing the cellular stress response. Full responsiveness to stress is only regained after 20 hours, at which time *PPP1R15A* mRNA returns to its basal level^18^. We therefore hypothesized that increased expression of GADD34 in DKO cells could reduce the responsiveness to stress and confer a higher level of stress adaptation. To test this hypothesis, we first assessed the stress responsiveness of DKO cell clones by measuring their ability to form SGs in response to increasing doses of sodium arsenite (Figures 7A and S8A). As hypothesized, the DKO clones showed reduced SG formation when exposed to intermediate doses of arsenite (100 to 250 µM), with the most striking difference at 100 µM, where approximately 50 % of the Ctrl cells formed SGs compared to only 20 % of the DKO cells. Importantly, the Huh7 GADD34ΔARE cells exhibited a similar reduction in responsiveness (Figures 7B and S8B), confirming that higher basal expression of GADD34 in Huh7 DKO and GADD34ΔARE cells attenuates their sensitivity to environmental stress, raising the threshold at which the ISR is triggered.

**Figure 7.**
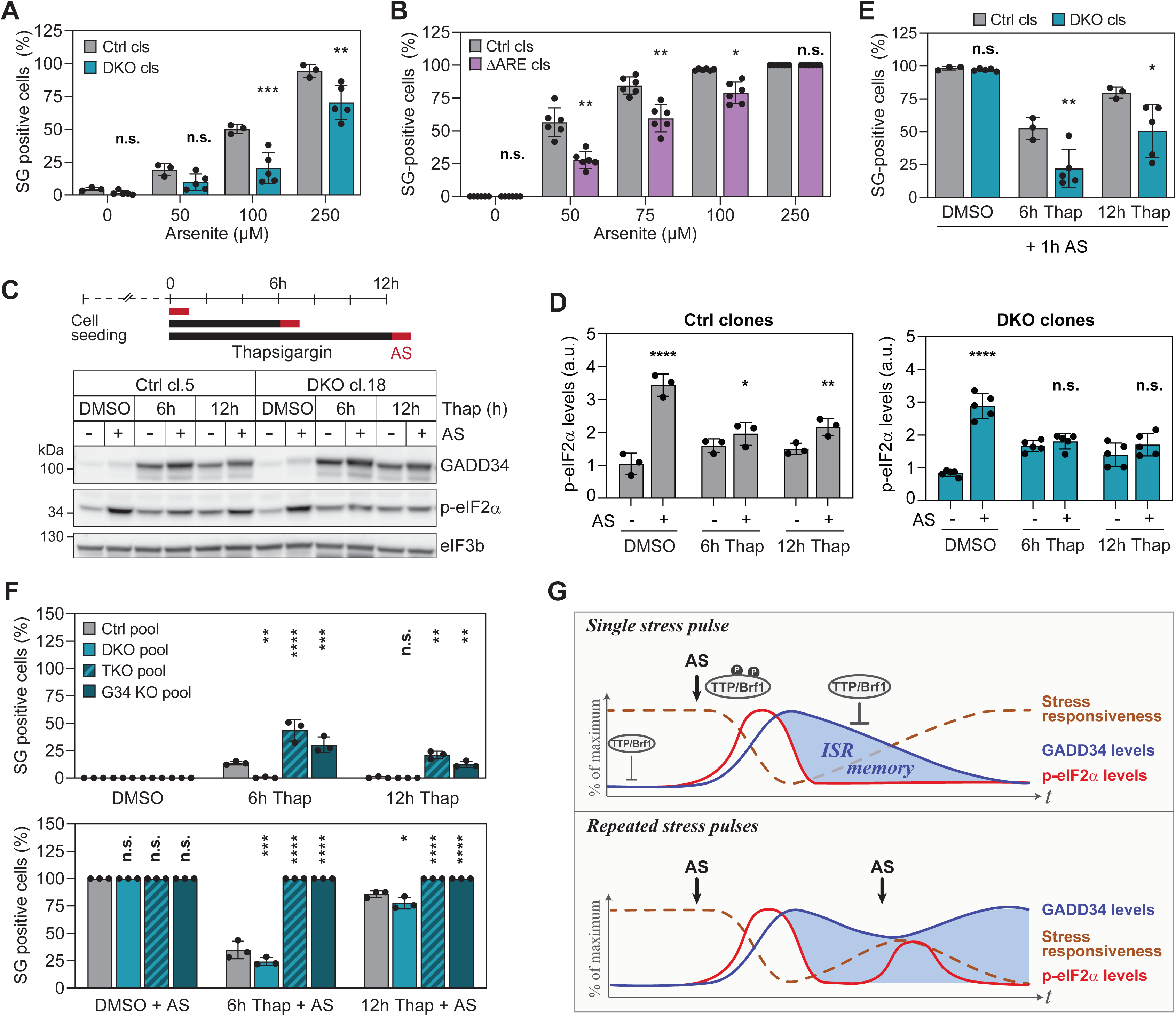
ZFP36 proteins contribute to set the threshold for stress responsiveness and the dynamics of stress adaptation. **(A and B)** Analysis of cell stress responsiveness to increasing doses of arsenite. The ability of cells to form SGs was assessed by immunofluorescence analysis. Shown is a representative mean percentage of SG-positive cells ± SD of all individual cell clones (n=2) for Huh7 Ctrl and DKO cell clones (A) and Huh7 Ctrl and GADD34ΔARE cell clones (B). Statistical significance compared to the respective Ctrl cell clones at the specified doses is indicated. **(C to E)** Huh7 Ctrl and DKO cell clones were treated with DMSO or thapsigargin for 6 or 12 h before a second stress pulse with 500 µM arsenite for 1 h (acute stress, AS). (C) Schematics of the experimental design (top panel) and representative Western blot analysis of Ctrl cl.5 and DKO cl.18 (lower panel). (D) Corresponding quantifications (mean ± SD) of p-eIF2α levels in all clones (n=2). Measured band intensities were normalized to eIF3b and depicted as arbitrary units (a.u.) (see also Figure S7D). Statistical significance compared to cells in the absence of AS (-) is indicated. (E) Analysis of SG formation in Huh7 Ctrl and DKO cell clones. Shown is a representative mean percentage of SG-positive cells ± SD of all individual clones (n=2). Statistical significance compared to Ctrl cell clones at the respective time point is indicated. **(F)** Role of GADD34 in response to on a second stress pulse applied during the stress adaptation phase. Huh7 Brf1/TTP/GADD34 triple knockout (TKO) and GADD34 knockout (G34 KO) cells were compared to Huh7 Ctrl and DKO cells. Cell pools were treated with DMSO or thapsigargin for 6 and 12 h and subsequently subjected to a second stress pulse of 500 µM arsenite for 1 h (acute stress, AS). Shown is the mean percentage of SG-positive cells ± SD of (n=3) prior to acute stress (top panel) and after acute stress (bottom panel). Statistical significance compared to Ctrl cell clones at the respective time point is indicated. **(G)** Proposed model: GADD34 levels act as a molecular memory of the ISR. Top panel: Exposure to an acute stress pulse results in phosphorylation of eIF2α and activation of the ISR, as well as inhibition of TTP and Brf1 decay activity by phosphorylation. Consequently, *PPP1R15A* mRNA synthesis is amplified by the stabilization of pre-existing and *de novo* synthesized mRNA, allowing a rapid increase in GADD34 levels and efficient dephosphorylation of eIF2α. Towards the stress adaptation phase, TTP and Brf1 are progressively dephosphorylated and regain their decay activity, destabilizing *PPP1R15A* mRNA. Elevated *PPP1R15A* mRNA levels act as a molecular memory of the previously activated ISR. Cells regain full responsiveness to stress once *PPP1R15A* mRNA turnover returns to its initial rate and levels return to baseline. Bottom panel: Cells subjected to a second acute stress pulse during the stress adaptation phase show attenuated responsiveness to stress due to *PPP1R15A* mRNA levels, which counteract eIF2α phosphorylation.

We next asked how prolonged expression of GADD34 during the stress adaptation phase would influence the responsiveness of DKO clones to a second stress stimulus. To this end, Huh7 Ctrl and DKO cell clones were pre-treated with thapsigargin for 6 or 12 hours, followed by an acute pulse of 500 µM arsenite, a dose at which 100% cells formed SGs (AS, Figure 7C). In agreement with our previous work^18^, pre-exposure of Ctrl clones to a 6-hour thapsigargin treatment strongly attenuated the phosphorylation of eIF2α (Figure 7D). This was associated with the formation of SGs in about 50 % of the cells (Figures 7E and S8C). In DKO cell clones, the pre-existing levels of GADD34 resulted in a more pronounced dephosphorylation of eIF2α, with only 20 % of cells forming SGs. At 12 hours, Ctrl cell clones had recovered almost entirely and 80 % of cells were able to form SGs, while DKO cell clones were still unable to mount an efficient SG response, with only about 50 % SG-positive cells (Figures 7E and S8C).

To confirm the impact of GADD34 on SGs formed in Huh7 Brf1/TTP DKO cells during repeated stress conditions, we used the ISR inhibitor ISRIB^62,63^, a small molecule that restores translation by antagonizing the inhibition of the guanine nucleotide exchange factor eIF2B by p-eIF2α. We confirmed ISRIB inhibitory activity in our cell model by co-treating Huh7 cells with a high, but non-cytotoxic, dose of ISRIB and thapsigargin or arsenite (Figures S8D, S8E, and S8F). ISRIB inhibited SG formation in approximately 75% of cells irrespective of the stressor concentration used (Figures S8E). We then pretreated Huh7 Ctrl and DKO cell pools with DMSO or thapsigargin for 6 and 12 hours prior to a 1-hour acute stress treatment with arsenite in the presence and absence of ISRIB. As shown in Figure S8G, depletion of TTP and Brf1 led to reduced SG formation under all conditions where cells were pretreated with thapsigargin as observed previously (Figure 7E). The reduction was most pronounced in the absence of pretreatment (DMSO) when ISRIB was administered together with the acute pulse of arsenite (Figure S8G), suggesting that TTP and Brf1 affect SG formation primarily via suppressing GADD34 expression.

To ascertain that GADD34 is a key target of TTP/Brf1 during stress adaptation, we generated Huh7 Brf1/TTP/GADD34 triple knock-out cells, referred to as TKO, as well as Huh7 GADD34 KO cells (Figures S9A and S9B). As anticipated, GADD34 depletion did not affect the ability of TKO and GADD34 KO cells to form SGs following a 1-hour exposure to thapsigargin (Figure S9C), yet it delayed SG clearance during the stress adaptation phase compared to Ctrl cells (Figures 7F top panel and S9C). In the absence of GADD34, it is likely that SGs are dissolved through the upregulation of ER-resident chaperones and/or other phosphatases such as CreP that can mediate eIF2α dephosphorylation. Under conditions of repeated stress, Huh7 TKO cells showed enhanced responsiveness to the second pulse of arsenite with 100% cells forming SGs despite thapsigargin pre-treatment, supporting the essential role of GADD34 during stress adaptation (Figure 7F bottom panel). These results highlight the importance of fine-tuning GADD34 levels during the adaptation phase and confirm that prolonged presence of GADD34 serves as a molecular memory for previous ISR activation. Together, these results provide compelling evidence that the post-transcriptional control of GADD34 expression by ZFP36 proteins is critical for controlling responsiveness and cellular adaptation to stress at the onset of the ISR and during the stress adaption phase.

## Discussion

Previous studies have suggested that *PPP1R15A* mRNA is intrinsically unstable^64^ and can be stabilized upon exposure to genotoxic stress^65^ and all-trans retinoic acid in a TTP-dependent manner^66^. However, the mechanism controlling *PPP1R15A* mRNA stability during the stress response remained elusive. In this work, we unveiled a novel component of ISR regulation at the level of the GADD34 negative feedback loop and identified control of ARE-mediated decay of *PPP1R15A* mRNA as a key mechanism that links the cellular experience of previous stress to its responsiveness towards novel stress exposure. Under basal conditions, recognition of the 3’ UTR ARE by members of the ZFP36 family results in the rapid decay of *PPP1R15A* mRNA, thereby attenuating GADD34 synthesis while ensuring its mRNA is continually produced. With an estimated half-life below 1.5 hours, *PPP1R15A* mRNA classifies among the five percent of fast-decaying cellular mRNAs^67^.

When cells encounter stress conditions, the regulatory circuit switches: the pre-existing pool of *PPP1R15A* mRNA is markedly stabilized within the acute phase of the stress response, due to phosphorylation-induced inactivation of ZFP36 proteins by the stress-activated p38-MAPK/MK2 and PI3K/Akt signaling pathways. The mechanism by which TTP and Brf1 are inactivated upon phosphorylation has been characterized previously: it involves binding of 14-3-3 adaptor proteins to phosphorylated TTP or Brf1^37,38^, which leads to reduced association with the deadenylase machinery that triggers mRNA decay^68^, and may further cause reduced affinity of ZFP36 proteins for ARE-containing target mRNAs^27^. This regulation promotes the timely production of GADD34 during the acute phase of the ISR, before *PPP1R15A* mRNA is transcriptionally and translationally induced as cells progress in the ISR. Stress-induced stabilization of *PPP1R15A* mRNA is transient and gradually decreases upon dephosphorylation and reactivation of ZFP36 proteins later during the stress adaptation phase. This mode of regulation resembles the stabilization of newly synthesized mRNAs during the acute phase of the unfolded protein response, in particular that of the stress induced *XBP1* mRNA, which escapes translational repression because of poly(A) tail extension^69,70^.

Our results provide mechanistic insight into how *PPP1R15A* mRNA levels are regulated. The persistence of *PPP1R15A* mRNA above basal levels as cells progress through the stress adaptation phase serves as a molecular memory of the previously activated ISR and allows for rapid translation of GADD34 in response to an increase in p-eIF2α levels when they encounter subsequent or continued stress. This molecular memory imposes a refractory phase and reduces responsiveness to stress by elevating the threshold at which the next ISR is triggered, thereby promoting cellular adaptation to acute and chronic stress of moderate intensity (Figure 7G).

The multilayer regulation of the GADD34 negative feedback loop underscores how important it is for cells to maintain low levels of *PPP1R15A* mRNA under basal conditions. In line with this notion, deletion of the ARE at the endogenous *PPP15R1A* locus led to a moderate, albeit significant, stabilization of *PPP1R15A* mRNA associated with enhanced translation and GADD34 expression. Notably, this effect was more pronounced during the stress adaptation phase. While a comparable phenotype has been previously reported upon deletion of the ARE in the TNF-α gene in mice^71^, other mechanisms may contribute to the rapid decay of *PPP1R15A* mRNA. For example, as reported for the yeast CPA1 mRNA, ribosome stalling at a uORF termination codon may activate nonsense-mediated mRNA decay^72^. Given that *PPP1R15A* mRNA translation in under control of two uORFs^9,10^, such a mechanism could potentiate the effect of ARE-mediated mRNA decay on *PPP1R15A* mRNA turnover. Other factors to consider are members of the ELAV family such as HuR, which have antagonistic functions to ZFP36 proteins and frequently stabilize ARE-containing mRNAs. Although HuR predominantly binds U- and CU-rich sequences, it recognizes over 80 % of the TTP binding sites^73^ and could therefore contribute to the stabilization of GADD34 mRNA under stress conditions. Depending on the cell type, differences in TTP, Brf1 and Hu expression levels may also play a role in determining the extent to which mRNA turnover controls GADD34 expression before, during and after the ISR.

Two of our observations suggest that ZFP36 family members are not simply redundant as temporal differences in their expression help to orchestrate the dynamics of GADD34 expression. First, their expression profile differs, with basal *ZFP36L1* mRNA levels encoding Brf1 being higher than those of *ZFP36* encoding TTP, while *ZFP36* mRNA expression levels exceed those of *ZFP36L1* in response to stress. Secondly, the results obtained with the single KO and DKO cell clones indicate that Brf1 activity is predominant under basal conditions, whereas TTP activity takes over in response to stress. Hence, it appears that the impact of ZFP36 family members follows the dynamics of their expression, and may render posttranscriptional control of *PPP1R15A* mRNA stability sensitive to different signaling pathways such as p38-MAPK (TTP and Brf1) and Akt (Brf1)^36–38^.

An important outcome of our study is that alterations in the post-transcriptional regulation of *PPP1R15A* mRNA and, consequently, GADD34 expression levels, critically affect cellular responsiveness and adaptation to stress. Defective ARE-mediated mRNA decay in TTP and Brf1 DKO cells as well as in GADD34ΔARE cells led to elevated basal expression of GADD34, which in turn reduced the responsiveness to stress. Following stress exposure, these cells required a longer period of time to regain full responsiveness. Comparison of Huh7 DKO, TKO and GADD34 KO cells showed that *PPP1R15A* mRNA is a key target of TTP/Brf1 for controlling stress responsiveness and adaptation. In addition to its essential role as an eIF2α phosphatase, GADD34 appears to have independent functions, e.g. in facilitating adaptation to osmotic stress by ensuring the preservation of Golgi integrity^74,75^. Increased GADD34 levels have also been reported to promote autophagy by suppressing mTOR signaling^76–78^, and both pro- and anti-apoptotic functions have been assigned to GADD34, depending on its expression level^79,80^.

By exploring the memory function of *PPP1R15A* mRNA in recording cellular stress experience, and demonstrating the impact of *PPP1R15A* mRNA stability on memory timing, our study uncovers an unexpected link between the regulation of mRNA turnover and the intricate translational control system that governs the ISR.

### Limitations of the study

While our study is based on human hepatocarcinoma cells and makes use of a wide range of stressors including viral infection, further work will be required to understand the importance of this mechanism systems that strongly depend on GADD34 such as pancreatic beta cells or in a mouse model subjected to physiologically relevant stresses. Furthermore, it is yet to be investigated whether Brf2, the third member of the ZFP36 family, contributes to this regulatory circuit. Future studies need to establish the role of GADD34 inherent instability to the tight temporal control of the negative feedback loop.

## Acknowledgements

We thank all members of the Ruggieri and Stoecklin laboratories for helpful discussions. We also thank the Infectious Diseases Imaging Platform (IDIP) at the CIID (Heidelberg University), and we are grateful to Christine Clayton at the Center for Molecular Biology of Heidelberg University (ZMBH) for providing access to the equipment required for polysome profile analysis. This work was supported by the Deutsche Forschungsgemeinschaft (DFG, German Research Foundation) [project numbers 240245660 SFB1129 TP13 to A.R., 278001972 TRR186 A14 to A.R. and G.S., 278001972 TRR186 A08 to V.H., 439669440 TRR319 RMaP A05 to A.R. and A04 to G.S., 464424253 CRC1550 to G.S., and 445549683 RTG2727 to J.S. and G.S.]; by the Biotechnology and Biological Sciences Research Council research grants [grant numbers BB/S006931/1 and BB/P068018/1 to N.L.].

## Author contributions

A.R. and G.S conceptualized the study. V.M., A.M., K.K., C.C.W., M.N., D.L., J.W., L.E. and J.S. performed and analyzed experiments. S.K. made a preliminary estimation of *PPP1R15A* mRNA stability using mathematical modelling. A.R., G.S., N.L. and V.M. drafted and edited the manuscript. V.H. critically revised the manuscript. All authors approved the final manuscript.

## STAR★Methods

### KEY RESOURCES TABLE

See additional Table

### RESOURCE AVAILABILITY

#### Lead Contact

Further information and requests for resources and reagents should be directed to and will be fulfilled by the Lead Contact, Alessia Ruggieri.

#### Materials Availability

All unique/stable reagents generated in this study are available from the Lead Contact with a completed Materials Transfer Agreement.

## METHODS

### Cell culture and culture conditions

Huh7 cells, HEK293T cells and Huh7 GADD34ΔARE, Huh7 TTP knockout (KO), Huh7 Brf1 KO, Brf1/TTP double knockout (DKO), Brf1/TTP/GADD34 triple knockout (TKO), and GADD34 KO cell clones were cultivated in complete Dulbecco’s modified Eagle medium (DMEM) supplemented with 10 % fetal calf serum (Capricorn), 2 mM L-glutamine, non-essential amino acids, 100 U/ml penicillin and 100 µg/ml streptomycin (all from Gibco). Huh7 cells stably expressing β-globin reporter constructs were generated by lentiviral transduction and subsequent selection in the presence of 5 µg/ml blasticidin (Blr).

### Generation of Huh7 knockout (KO) cell clones

A two-crRNA strategy was used for the deletion and/or knockout of genomic sequences of interest. The crRNAs (pre-designed or custom-designed) were purchased from Integrated DNA technologies (IDT).

For the deletion of the ARE sequence from the endogenous *PPP1R15A* locus, two custom-designed crRNAs flanking the ARE sequence: sequence 1 (5’ – / AltR1 / GAG GCG UGG CUG AGA CCA ACG UUU UAG AGC UAU GCU / AltR2 / – 3’) and sequence 2 (5’– / AltR1 / UAU UUA UUU UUU CUA AGU GGU UUU AGA GCU AUG CU / AltR2 / – 3’).

For the knockout of *ZFP36* (TTP) and *ZFP36L1* (Brf1) genes: two pre-designed crRNAs per target gene: Hs.Cas9.ZFP36.1.AA (5’ – / AltR1 / CCG UGC CAU CCG ACC AUG GAG UUU UAG AGC UAU GCU / AltR2 / – 3’) and Hs.Cas9.ZFP36.1.AC (5’ – / AltR1 / CCA CAA CCC UAG CGA AGA CCG UUU UAG AGC UAU GCU / AltR2 / – 3’), Hs.Cas9.ZFP36L1.1.AB (5’ – / AltR1 / GGG UGA CUG AGU GCC UCC GAG UUU UAG AGC UAU GCU / AltR2 / – 3’) and Hs.Cas9.ZFP36L1.1.AR (5’ – / AltR1 / UGC CGC ACC UUC CAC ACC AUG UUU UAG AGC UAU GCU / AltR2 / – 3’).

For the knockout of *PPP1R15A* (GADD34): Hs.Cas9.PPP1R15A.AC (5’– / AltR1 / AUC ACU GGG UUG GCA CCA CCG UUU UAG AGC UAU GCU / AltR2 – 3’) and custom-designed (5’ - AltR1 / GAG GCG UGG CUG AGA CCA ACG UUU UAG AGC UAU GCU / AltR2 – 3’).

A negative control crRNA (Integrated DNA technologies) was used to generate control cell lines. Individual crRNAs and tracrRNA (Integrated DNA technologies) were resuspended in 1 x TE buffer (10 mM Tris, 0.1 mM EDTA, pH 7.5) to a final concentration of 200 µM and hybridized at equimolar amounts by heating to 95 °C for 5 min and slow stepwise cooling to room temperature. Pre-assembled crRNA/Cas9 ribonucleoprotein complexes (RNPs) were obtained by gently mixing 1.2 µl of hybridized RNAs with 17 µg of recombinant Cas9 protein (Integrated DNA technologies) and 2.1 µl of PBS, followed by 20 min incubation at room temperature.

Huh7 cells (2 x 10^6^) were washed with PBS, resuspended in 91 µl supplemented Solution T (Lonza) and mixed with 2.5 µl of each pre-assembled RNP and 4 µl electroporation enhancer (Integrated DNA technologies). Nucleofection was performed using an Amaxa 2b nucleofector (Lonza) and pre-set program T-22. Nucleofected cells were incubated for 48 h prior to clonal amplification and screening for homozygous clones using target-specific PCR (GoTaq Hot Start, Promega) on genomic DNA using the following primers: TTP-KO-For: 5’–ACC TCC AAC TCT GGG TTC CT–3’; TTP-KO-Rev: 5’–GAG AAG GCA GAG GGT GAC AG–3’; Brf1-KO-For: 5’–GTG GTA GCT TTG GGG ACA GA–3’; Brf1-KO-Rev: 5’–GTC ATC GGC GCT CAG AAT AG–3’; ARE-seq-long-For 5’–ACC TTG CCT TCC TCC TCT GTC CCT–3’; ARE-seq-long-Rev: 5’–GCT ATT CTC TTC CCA GCT CCC TCA–3’. GADD34-KO-For: 5’–ATC ACA CCG GGA GTG TTG TC –3’; GADD34-KO-Rev: 5’–AGA CGC TGC TCG CTA CAA AT–3’.

For Huh7 GADD34ΔARE cell clones, the resulting PCR product was purified and directly send for sequencing. For TTP KO, BRF1 KO, Brf1/TTP DKO, Brf1/ TTP/GADD34 TKO and GADD34 KO cell clones, PCR product pools were further subcloned into a TOPO TA cloning vector (Thermo Fisher Scientific) according to manufacturer’s recommendation and transformed into DH5α competent bacteria. Eight to ten colonies were selected and amplified for the extraction of plasmid DNA, which was then send for sequencing to ensure that all potential allelic sequence variants were covered.

### Plasmids

Plasmids encoding mouse ZFP36 proteins, i.e. pcDNA3-TTP-mycHis (mTTP), pcDNA3-TTP-m1,2-mycHis (mTTP_1,2_), and pcDNA3-TTP-AA-mycHis (mTTP-AA) have been described elsewhere^37,81^.

Plasmids encoding human ZFP36 proteins TTP and Brf1: human *ZFP36* and *ZFP36L1* open reading frames were amplified from the ORFeome cDNA clone library (Invitrogen, Life Technologies) by Phusion PCR (Thermo Fisher Scientific) using primers flanked by ClaI and XbaI restriction enzymes (New England Biolabs), digested and inserted into pcDNA3-TTP-mycHis vector backbone digested with the same restriction enzymes. ClaI-huTTP-For: 5’–ATA ACT ATC GAT ATG GAT CTG ACT GCC ATC TAC GAG– 3’; XbaI-huTTP-Rev: 5’–ATT ACC TCT AGA CTC AGA AAC AGA GAT GCG ATT GAA–3’; ClaI-huBrf1-For: 5’–ATA TTT ATC GAT ATG ACC ACC ACC CTC GTG TCT GCC–3’; XbaI-huBrf1-Rev: 5’–GGC GAC TCT AGA GTC ATC TGA GAT GGA AAG TCT GCT–3’. For transient expression, sequences of all myc-his tagged ZFP36 proteins were digested using ClaI and PmeI restrictions enzymes (New England Biolabs), blunted using DNA polymerase I large Klenow fragment (New England Biolabs) and inserted into the lentiviral vector pWPI Puro carrying the puromycin resistance gene, linearized with PmeI.

Plasmids pcDNA3-7B β-globin reporter (WT) and pcDNA3-7B TNFα ARE containing the ARE of mouse TNF-α (mTNFα-ARE) were described eslewhere^82,83^. β-globin reporter constructs expressing GADD34 3’UTR (pcDNA3-7B-β-globin reporter GADD34 human 3’UTR) or GADD34 3’UTR with mutated ARE pentamer motives (pcDNA3-7B-β-globin reporter GADD34 human mutated 3’UTR) were generated by insertion of the corresponding sequences upstream of the β-globin 3’ UTR sequence using hybridized oligonucleotides with compatible overhangs to those produced by BglII digest. The following oligonucleotides were used: GADD34-3’UTR-For: 5‘–GAT CTG ACC AAC TGG TTT GCC TAT AAT TTA TTA ACT ATT TAT TTT TTC TAA GTG TGG GTT TAT ATA AGG A–3’ and GADD34-3’UTR-Rev: 5‘–GAT CTC CTT ATA TAA ACC CAC ACT TAG AAA AAA TAA ATA GTT AAT AAA TTA TAG GCA AAC CAG TTG GTC A–3’; GADD34-3’UTR-mut-For: 5‘–GAT CTG ACC AAC TGG TTT GCC TAT AAT GTA TTA ACT ATG TAT TTT TTC TAA GTG TGG GTT TAT ATA AGG A–3‘ and GADD34-3’UTR-mut-Rev: 5‘–GAT CTC CTT ATA TAA ACC CAC ACT TAG AAA AAA TAC ATA GTT AAT ACA TTA TAG GCA AAC CAG TTG GTC A–3‘. Equimolar amounts of complementary oligonucleotides were phosphorylated by incubation with T4 polynucleotide kinase in T4 ligation buffer (New England Biolabs) for 30 min at 37°C. Phosphorylated oligonucleotides were hybridized by heating the reaction to 95°C for 5 min followed by slow stepwise cooling to room temperature. Hybridized oligonucleotides were then inserted into pcDNA3-7B digested with BglII. For the production of Huh7 stable cell lines, all β-globin reporter sequences were were excised from the pcDNA3-7B constructs using HindIII and XbaI, blunted using DNA polymerase I large Klenow fragment and inserted into the lentiviral vector pWPI Blr, carrying the blasticidine resistance gene^84^, linearized with PmeI (pWPI -β-globin GADD34 3’UTR Blr and pWPI -β-globin GADD34 3’UTR mut Blr).

### Lentivirus production, transient transduction and stable cell lines

HEK293T cells (5 x 10^6^) were seeded on the day prior to transfection. Cells were transfected with 6.42 µg pWPI lentiviral vector, 6.42 µg packaging vector pCMV Δ8.91 and 2.16 µg pMD2.G encoding VSV-G glycoprotein (kind gift from Didier Trono)^85^. DNA was diluted with OptiMEM (Gibco) and mixed with polyethylenimine (PEI) solution (pH7.2) in a 1:3 (PEI:DNA) ratio. Samples were vortexed vigorously prior to incubation at room temperature for 20 min. Transfection mix was added dropwise to the cells and transfected cells were incubated for 48 h. Medium was replenished at 6 h post-transfection. The lentivirus-containing supernatant was filtered through 0.45 µm pore-size filters and stored at -80°C or directly used to transduce target Huh7 cells. For the generation of stable cell lines, the corresponding antibiotic selection was added for 36 h after transduction. For transient transductions, Huh7 were directly seeded into lentiviral supernatant diluted 1:10 in complete DMEM with the following cell densities: 8 x 10^4^ cells for RNA decay experiments or 2.5 x 10^5^ cells for protein analysis and polysome profiling. Medium was replenished after 24 h and cells used for further analysis at 40 h post transduction.

### Virus infections

Huh7 cells were infected with hepatitis C virus (HCV Jc1) at an MOI of 5 for 24 h or with trans-complemented HCV particles (HCV_TCP_) at an MOI of 10 TCID_50_ for 48 h prior to the addition of 100 U/ml IFN-α for 24 h to induce an antiviral stress response as previously described^17,18^. Alternatively, Huh7 cells were infected with dengue virus (DENV) serotype 2 strain New Guinea C (kindly provided by Andrew Davidson) at an MOI of 10 for 8 h or 30 h prior to actinomycin D chase.

### Transfection with double-stranded (ds) RNA

Synthesis and purification of 200-bp dsRNA was described previously^18^. For *PPP1AR15A* mRNA turnover analyses, 1.7 x 10^5^ Huh7 cells were seeded in a 6-well plate and transfected for 16 h with 1 µg purified dsRNA mixed with Lipofectamin 2000 transfection reagent (Thermo Fisher Scientific) in a 1:3 ratio prior to actinomycin D chase.

### Chemical stress treatments

Huh7 cells were seeded one day prior to treatment (5 x 10^4^ cells for RNA extraction and 3 x 10^5^ cells for protein analyses). Thapsigargin (Biotrend) was dissolved in DMSO and used at a concentration of 2 µM for up to 12 h. Sodium arsenite (Sigma Aldrich) was resuspended in H_2_O and used at a concentration of 500 µM for 45 min for the induction of acute stress response and at the indicated increasing concentrations for stress responsiveness experiments. ISRIB was resuspended in DMSO (Sigma Aldrich) and used in co-treatment with 500 µM arsenite at a concentration of 1000 nM for 1 h.

### Cell viability assay

Huh7 cells were seeded in triplicate wells (2 x 10^4^ cells) one day prior to treatment. Cell viability was measured using the Cell Proliferation Reagent WST-1 (Roche) as recommended by the manufacturer. After 1 h incubation with increasing concentrations of ISRIB, fluorescence was measured at 440 nm with a reference wavelength >600 nm using a Tecan microplate reader. Values were normalized to the DMSO control and depicted as percentages.

### Quantitative PCR Analysis

All samples were analyzed in triplicates using a CFX96 Touch Real-Time PCR detection system (Bio-Rad). For probe-based RT-qPCR, 3 µl of extracted total RNA were mixed 15 µl qPCR mix (qPCRBIO Probe 1-Step Go Lo-ROX, PCR Biosystems) according to manufacturer’s protocol. For the analysis of *PPP1R15A*, *GAPDH* and *firefly* luciferase transcript relative abundance, absolute RNA copy numbers were calculated with the help of a standard curve (purified *in vitro* transcripts). The following cycling protocol was used: 50°C 10 min, 95 °C 1 min, 95°C 10 sec, 60°C 1 min; steps 3 – 4 repeated for 39 times. The following primers and probes were used: GADD34-For: 5’–CAG AAA CCC CTA CTC ATG ATC C–3’; GADD34-Rev: 5’–AAA TGG ACA GTG ACC TTC TCG–3’; GADD34-pre-mRNA-For: 5’–ACA GTG ACA GGC AAG TGA CTA G–3’; GADD34-Pre-mRNA-Rev: 5’–GGA AGA GAG AGA GAG AAG CAA AC–3’; GADD34-Probe-FAM: 5’–6–FAM–CCC CTA AAG GCC AGA AAG GTG CGC–TAMRA–3’; GAPDH-For: 5’–GAA GGT GAA GGT CGG AGT C–3’; GAPDH-Rev: 5’–GAA GAT GGT GAT GGG ATT TC–3’; GAPDH-Probe-VIC: 5’–VIC-CAA GCT TCC CGT TCT CAG CCT-TAMRA–3’; *firefly*-luciferase-For: 5’–CCC TGG TTC CTG GAA CAA TT–3’; *firefly*-luciferase-Rev: 5’–ATA GCT TCT GCC AAC CGA AC–3’; *firefly*-luciferase-Probe-Cy5: 5’– Cy5-ATC GAG GTG GAC ATC ACT TAC GCT-BHQ3–3’; β-globin-For: 5’–GAA GGC TCA TGG CAA GAA GG–3’; β-globin-Rev: 5’–ATG ATG AGA CAG CAC AAT AAC CAG–3’; β-globin-Probe-FAM: 5’–6-FAM-ACA AGC TGC ACG TGG ATC CTG AGA A-BHQ1–3’.

For two step SYBR-green based RT-qPCR, total RNA was first reverse transcribed using the High-Capacity cDNA Reverse Transcription kit as recommended by the manufacturer (Applied Biosystems, 25°C 10 min, 37°C 2 h, 85°C 5 min). The resulting cDNA was diluted 1:20 in RNase-free H_2_O and 3 µl dilution were used for RT-qPCR analysis using iTaq Universal SYBR Green Supermix (Bio-Rad) according to manufacturer’s instructions and using the following cycling protocol: 95°C 3 min, 95°C 10 sec, 60°C 30 sec – with step 3-4 repeated for 45 times. *GAPDH* mRNA values were used for normalization. Changes in expression levels were calculated using the ΔCT method and expressed as arbitrary units (a.u.). Fold changes in expression were calculated using the ΔΔCT method^86^.

### Measurements of mRNA turnover

Huh7 cells (5 x 10^4^ cells) were treated with 5 μg/ml of actinomycin D in DMSO (Sigma-Aldrich). Samples were harvested at the indicated times for total RNA extraction using the NucleoSpin RNA kit (Macherey Nagel) following the manufacturer’s protocol, and further analyzed by probe-based qPCR. For the absolute quantification of *PPP1R15A* and *GAPDH* mRNA half-lives shown in Figure 1A, the number of *PPP1R15A* and *GAPDH* mRNA copies at each time point was normalized to the copy numbers of spiked *firefly* luciferase transcripts spiked into the sample to account for RNA loss during purification. As *GAPDH* mRNA levels did not significantly change over the course of the experiment, the relative abundance of the *PPP1R15A* mRNA was normalized to that of *GAPDH* mRNA in all subsequent calculation of mRNA half-life panels using the ΔCT method. Fold changes in abundance relative to T0 were calculated using the ΔΔCT method^86^ and expressed as a percentage. The resulting decay curves were obtained by using a non-linear regression (GraphPad Prism) to estimate the mRNA half-life (t_1/2_). For half-lives exceeding 10 h or which could not be determined, values are indicated as “n.d. (> 10 h)”.

For the analysis of the turnover of the newly synthesized pool of *PPP1R15A* mRNA, 5 x 10^6^ Huh7 cells were seeded in 15-cm dishes the day prior to the experiment. Cells were incubated with medium containing either 0.2 mM 5-EU analog and 2 µM thasigargin or with 0.2 mM 5-EU analog and an equal volume of DMSO, for 2 h at 37°C. Next, cells were detached with trypsin, pelleted by centrifugation, resuspended in 20 ml of DMEM containing 10 µg/ml of actinomycin D and equally distributed into five 14-ml Falcon tubes. Cells were incubated for 10 min at 37°C before starting the 2-hour chase experiment. At each time point, cells were harvested by pelleting and lysed by addition of 750 µl TRIzol reagent (Invitrogen). Total RNA was extracted as recommended by the manufacturer. The newly synthesized RNA, which had incorporated the 5-EU analog, was further isolated using the Click-iT Nascent RNA Capture Kit (Invitrogen), according to the manufacturer instructions. Briefly, for each time point, 5 µg of total RNA was biotinylated with an azide-modified biotin in a copper catalyzed click reaction. Next, 500 ng of biotinylated total RNA was used for subsequent capture with 50 µl of streptavidin magnetic beads. After several wash steps, the bead suspension was heated at 68–70°C for 5 min and the biotinylated RNA captured on the beads immediately used as a template for cDNA and SYBR Green-based qPCR (see quantitative real-time PCR (RT-qPCR) analysis). Because of the different transcription rate of the housekeeping genes, *PPP1R15A* mRNA levels were normalized to the value of each time point to that of T0.

For the analysis of the turnover of the pre-existing pool of *PPP1R15A* mRNA, 2 x10^5^ Huh7 cells were treated with 10 µg/ml of actinomycin D and incubated for 10 min at 37°C to block transcription. Control cells were treated with DMSO to measure *PPP1R15A* mRNA transcriptional induction during the subsequent thapsigargin treatment. Transcriptional induction of *PPP1R15A* mRNA in response to thapsigargin treatment was analyzed by measuring the levels of *PPP1R15A* pre-mRNA. To this end, 30 µl of total RNA were incubated with 1 µl TURBO DNAse and TURBO DNase buffer (Invitrogen) in a total volume of 50 µl and incubated for 30 min at 37°C to remove residual genomic DNA. The DNAse-digested RNA was purified using the Monarch RNA Cleanup kit (New England Biolabs) as recommended by the manufacturer and used as template for cDNA synthesis and SYBR Green-based qPCR (see quantitative real-time PCR (RT-qPCR) analysis). After transcription inhibition, cells were treated with 2 µM of thapsigargin or DMSO for 1 h at 37°C prior to the 2-hour actinomycin D chase. At each time point, cells were harvested by addition of 750 µl of TRIzol reagent and RNA extracted as recommended by the manufacturer. *PPP1R15A* mRNA levels were normalized to that of *GAPDH* mRNA using the ΔCT method. Fold changes in abundance relative to T0 were calculated using the ΔΔCT method.

### Transcriptome-wide mRNA half-life measurements

Prior to rRNA depletion using the NEBNext rRNA Depletion Kit v2 (New England Biolabs), 1 μg total RNA was mixed with 2 μl 1:100 diluted ERCC RNA Spike-In Mix (Thermo Fisher Scientific). Libraries were prepared with the NEBNext Ultra II Directional RNA Kit (NEB). Samples were equimolarly pooled and sequenced on a NextSeq550 system (Illumina) with 80 bases single-end. Reads were mapped to the human genome (hg38, using Gencode V27 as downloaded from the UCSC Genome browser wgEncodeGencodeBasic27 table as transcriptome annotation) and the ERCC92 spike-in mix with STAR v2.5.3a^87^, allowing up to 2 mismatches. Read counts were summarized at the gene level with the featureCounts function of the subread package v1.6.3^88^. A read was only counted when it was completely contained within an exon. Read counts were normalized to library size using the median of ratios method of the DESeq2 package^89^.

### RNA immunoprecipitation

Transduced Huh7 cells (1.6 x 10^6^ cells) were washed twice with ice-cold PBS and lysed by scraping cells on ice with 1 ml IP lysis buffer (50 mM Tris HCl pH 8.0, 150 mM NaCl, 1 mM MgCl_2_, 1 % NP40, 2 % glycerol, 1 mM DTT, 1 mM Na_3_VO_4_, 50 mM NaF, supplemented with cOmplete EDTA-free protease inhibitor (Roche)). Lysates were incubated for 10 min on ice and subsequently centrifuged at 20,800 x g for 10 min at 4 °C. Supernatants were transferred into DNA LoBind tubes (Eppendorf) and two aliquots of 40 µl each were kept for later RNA and protein input analyses. Binding conditions were optimized by adding NaCl and imidazole to the lysates to achieve a final concentration of 250 mM NaCl, 10 mM imidazole. Ni-NTA agarose beads (Qiagen) were pre-washed twice with 500 µl IP lysis buffer and added to samples (50 µl/sample). After tumbling at 4 °C for 2 h, beads were washed 6 times by the addition of 1 ml IP wash buffer (50 mM Tris HCl pH 8.0, 250 mM NaCl, 1 mM MgCl_2_, 1 % NP40, 2 % glycerol, 1 mM DTT, 1 mM Na_3_VO_4_, 50 mM NaF, 10mM imidazole, supplemented with protease inhibitor) and centrifugation at 2,650 x g for 3 min at 4 °C. After the last wash, samples were split, with one half used for protein elution and the other half for RNA extraction. For protein elution, beads were centrifuged once at 5,000 x g and resuspended in 30 µl IP elution buffer (50 mM Tris HCl pH 8.0, 500 mM NaCl, 1 mM MgCl_2_, 1 % NP40, 2 % glycerol, 1 mM DTT, 1 mM Na_3_VO_4_, 50 mM NaF, 500 mM imidazole, supplemented with protease inhibitor). After 2 min incubation on ice, samples were centrifuged, and supernatants transferred into fresh tubes. The elution procedure was repeated once and both eluates were pooled and stored at -80°C until further analysis. Prior to RNA extraction, 360 µl of IP lysis buffer were added to RNA input samples to achieve equal sample volumes. 400 µl RNA extraction buffer (10 mM Tris HCl pH 7.5, 350 mM NaCl, 10 mM EDTA, 1 % SDS, 42 % urea) was added to lysates and each sample was spiked with 3 x 10^8^ copies of *firefly* luciferase *in vitro* transcripts to allow for later normalization of the values to RNA loss during purification. After the addition of 400 µl phenol/chloroform/isoamyl alcohol (25:24:1, v/v, Applichem) samples were vigorously vortexed prior to centrifugation at 20,800 x g for 15 min at room temperature. The aqueous upper phase was carefully recovered and transferred to fresh tubes. RNA was precipitated by addition of 800 µl isopropanol containing 0.2 % GlycoBlue (Invitrogen) and samples were inverted several times prior to overnight incubation at - 20 °C. RNA was pelleted by centrifugation at 20,800 x g for 15 min at 4 °C, washed once with 70 % ethanol and shortly air-dried prior to resuspension in 40 µl RNAse-free water. Absolute abundance of *PPP1R15A* or β-globin mRNA was further analyzed by probe-based qPCR. RNA enrichment following protein-RNA co-IP was calculated as follows: CT values of *PPP1R15A* or β-globin reporter mRNAs were log transformed (1/2^CT^) and total RNA copies of spiked *firefly* luciferase *in vitro* transcript were used to account for RNA loss during purification. To account for differences in IP efficiency, values from the precipitated fractions (IP) were further normalized to the absolute protein signal intensities measured in the precipitated samples. Normalized values are depicted as IP/Input and fold GFP control.

### SDS-PAGE and Western blot analysis

Cells were washed with ice-cold PBS and lysed by scraping in ice-cold protein lysis buffer (50 mM Tris HCl pH 7.4, 150 mM NaCl, 1 % Triton X-100, 60 mM β-glycerophosphate, 15 mM 4-nitrophenylphosphate, 1 mM Na_3_VO_4_, 1 mM NaF, supplemented with cOmplete EDTA-free protease inhibitor (Roche)). Cell lysates were incubated on ice for 30 min and subsequently centrifuged at 20,800 x g, 4 °C for 15 min. Supernatants were transferred in fresh tubes and stored at -80 °C until analysis. Protein concentration was determined using the Protein Assay Dye Reagent Concentrate (Bio-Rad). Forty µg (for the analysis of p-eIF2α, Akt, and p38) or 100 μg (for the analysis of GADD34, p-Akt, and p-p38) total protein was mixed with 6x SDS sample buffer (375 mM Tris-HCl pH 6.8, 50 % glycerol, 9 % SDS, 9 % β-mercaptoethanol, 0.03 % bromophenol blue) and separated by standard polyacrylamide gel electrophoresis. Proteins were transferred onto PVDF membranes (Millipore). Membranes were blocked in 5 % protease-free BSA (Sigma Aldrich) or 5 % non-fat dry milk diluted in TBS-T buffer (25 mM Tris-HCl, 150 mM NaCl, 2 mM KCl, pH 7.4, 0.1 % Tween), depending on antibody requirements. The following primary antibodies were used at the indicated dilutions and incubated overnight at 4 °C : anti-GADD34 (Proteintech, Cat# 10449-I-AP; 1:1,000; milk), anti-eIF2α (Cell Signaling, Cat# 9722; 1:1,000; milk), anti-phospho-eIF2α (Cell Signaling, Cat# 3398; 1:1,000; BSA), anti-p38 (Cell Signaling, Cat# 8690; 1:1,000 milk), anti-phospho-p38 (Cell Signaling, 4511, 1:1,000; BSA), anti-Akt (Cell Signaling, Cat# 4691; 1:1,000; milk), anti-phospho-Akt (R&D systems, Cat# AF887; 1:1,000; BSA), anti-HCV NS5A (NS5A 9E10 was a gift from Charles Rice; 1:2,000; milk), anti-GAPDH (Santa Cruz, Cat# sc-365062; 1:5,000; milk), anti-eIF3a (Bethyl Laboratories, Cat# A302-002A; 1:5,000; milk), anti-eIF3b (Bethyl Laboratories, Cat# A301-761A; 1:5,000; milk), anti-myc-tag (Cell Signaling, Cat# 2276; 1:1,000; milk), anti-puromycin 12D10 (Millipore, Cat# MABE343; 1:2000; milk), anti-Tristetraprolin (TTP) D1I3T (Cell Signaling, Cat# 71632; 1:500; milk), anti-ZFP36L1 (Brf1) (Thermo Fisher Scientific, Cat# PA5-68973; 1:500; milk). Membranes were washed thrice with TBS-T buffer prior to addition of horseradish peroxidase (HRP)-conjugated secondary antibodies diluted in blocking solution (anti-mouse-HRP 1:10,000 or anti-rabbit-HRP 1:10,000 antibodies; all from Sigma Aldrich) for 1 h at room temperature. Membranes were washed 3 times for 5 min in TBS-T buffer and prior to chemiluminescence detection using Western Lightning chemiluminescent substrate (Perkin Elmer). Chemiluminescence signal was detected with an ECL-Imager (INTAS) and band quantification performed using the LabImage1D software (INTAS). The measured band intensities were normalized to the indicated loading controls and are shown as arbitrary units (a.u.) or fold control. To account for the variation in total signal intensity among repeats, biological repeats that were imaged separately were additionally normalized to the sum of signals from bands of this membrane^90^.

### Phos-tag acrylamide gel analysis

For Phos-tag gels, 8 % acrylamide gels were prepared by adding 70 µM Phos-tag (FUJIFILM Wako Chemicals) and 140 µM MnCl_2_ to the resolving gel before casting. For the preparation of dephosphorylated protein samples, cells were lysed in dephosphorylation buffer (50 mM Tris HCl pH 8.0, 150 mM NaCl, 1 mM MgCl_2_, 1 mM MnCl_2,_ 1 % NP40, 2 % glycerol, supplemented with cOmplete EDTA-free protease inhibitor (Roche)). 100 µg cell lysate was mixed with 3,200 U λ protein phosphatase (New England Biolabs) and incubated at 37 °C for 1 h to achieve complete protein dephosphorylation. 100 µg of cell lysate or dephosphorylated lysates were separated on the Phos-tag acrylamide gels and ran in TGS running buffer (25mM Tris-HCl pH 8.8, 192 mM glycine, 0.1% SDS in dH_2_O) for 2 h in the dark. After electrophoresis, gels were washed in wet blot transfer buffer (25 mM Tris Base, 150 mM Glycine, 20 % (v/v) Methanol, pH 8.3) supplemented with 1 mM EDTA for 10 min, followed by an additional washing step in absence of EDTA. Proteins were then transferred on pre-activated PVDF membranes by wet blot transfer. Membranes were blocked, stained with the anti-myc-tag antibody, and imaged as described above. For quantification, the signal from the upper, hyper-phosphorylated band as well as the total lane signal were quantified, and the ratio of hyper-phosphorylated/total protein was calculated.

### Polysome profiling

Huh7 cells (1 x 10^6^) were seeded one day prior to the experiment. Translation was arrested by treatment with 100 µg/ml cycloheximide (Sigma Aldrich) for 10 min at room temperature. Cells were washed with ice-cold PBS supplemented with 100 µg/ml cycloheximide and lysed by scraping in polysome lysis buffer (20 mM Tris-HCl pH 7.4, 5 mM MgCl_2_, 150 mM NaCl, 1 % Triton X-100, 100 µg/ml cycloheximide, 200 U/ml RNAsin, 1 µl/ml β-mercaptoethanol; supplemented with cOmplete EDTA-free protease inhibitor cocktail (Roche)). Cell lysates were tumbled for 10 min at 4°C and nuclei pelleted by centrifugation at 10,000 x g for 10 min at 4°C. Cell lysate was loaded onto a linear 17.5–50 % sucrose gradient (dissolved in 20 mM Tris-HCl pH 7.5, 5 mM MgCl_2_, 150 mM NaCl) and separated by ultracentrifugation at 35,000 rpm in a SW60 rotor (Beckman) at 4 °C for 2.5 h. Polysome profiles were recorded by measuring the absorbance at 254 nm using Teledyne ISCO Foxy R1 system in combination with PeakTrak software. Fractions of approximately 300 μl were collected every 14 seconds. For RNA isolation, fractions were eluted in 300 µl urea buffer (10 mM Tris pH 7.5, 350 mM NaCl, 10 mM EDTA, 1 % SDS and 7 M urea) and spiked with 3 x 10^8^ copies of *firefly* luciferase *in vitro* transcript to account for loss of material during purification. Total RNA was purified by phenol/chloroform/isoamyl alcohol (25:24:1, v/v, Applichem) precipitation as described above. RNA samples were analyzed on a 1 % agarose gel to distinguish monosomal and polysomal fractions. Abundance of *PPP1R15A* mRNA, *GAPDH* mRNA, β-globin reporter mRNA and *firefly* luciferase *in vitro* transcript was measured in each fraction using RT-qPCR as described above. CT values of target RNAs in individual fractions were log transformed (1/2^CT^) and normalized to the *firefly* luciferase spike-in control. For the analysis of individual fractions, normalized values are depicted as percentages of the sum of total RNA measured in all fractions (cumulative mRNA distribution). The proportion of polysome association was estimated by calculating the area above the cumulative mRNA distribution and statistical significance calculated using one-way ANOVA.

### Immunofluorescence microscopy

Cells were seeded onto glass coverslips one day before drug treatment, washed once with PBS and fixed for 15 min fixed with 4 % paraformaldehyde (PFA) in PBS. Cells were permeabilized with 0.5 % Triton-X in PBS for 5 min, washed with PBS and blocked for 30 min in blocking buffer (5 % horse serum, 5 % sucrose in PBS) to reduce unspecific binding. Antibodies were diluted in blocking buffer and added for 1 h at the following dilutions: anti-eIF3η (Santa Cruz, Cat# sc-16377 and Cat# sc-137214, both 1:500). Cells were washed 3 times for 5 min with PBS. Secondary antibodies (donkey anti-mouse Alexa Fluor 488, donkey anti-goat Alexa Fluor 568; all from Molecular Probes) were diluted 1:2,000 in blocking buffer and used for detection of primary antibodies for 1 h at room temperature. Nuclei were stained with 25 ng/ml DAPI (Roche) in PBS, for 3 min prior to washing. Cells were washed twice with PBS and once with H_2_O. Coverslips were mounted on glass slides using Fluoromount-G mounting medium (SouthernBiotech). Images were acquired on a Nikon Eclipse Ti-E microscope using 20x (N.A. 1.0 PLAN APO λ) or 40x (N.A. 1.3 PLAN Fluor Oil) objectives. Image analysis was performed using the FIJI software package^91^.

For SG disassembly experiments, Huh7 cells were seeded on 6-well plates containing glass coverslips the day prior the experiment and treated with thapsigargin for 1 h before removal of the drug, washing 3 times with sterile PBS and adding fresh complete DMEM. Cells were further incubated at 37°C, harvested and fixed at the indicated time-points post drug removal using 4% PFA in PBS.

### Ribopuromycylation assay

*De novo* synthesized proteins were quantified by measuring the incorporation of puromycin into native peptide chains described previously ^59,92^ and using Western blot analysis. Huh7 cells (2×10^5^) were treated with DMSO or thapsigargin for the indicated time points. Control cells were treated with 200 µg/ml cycloheximide for 1 h to inhibit host cell translation elongation. Shortly before harvesting, cells were incubated with 10 µg/ml puromycin (Gibco) for 5 min at 37°C, washed twice with PBS, and harvested for SDS-PAGE and Western blot analysis as described above. Before blotting, membrane was stained using a Ponceau solution (0.01% Ponceau S in 1% acetic acid) for total protein normalization. Released puromycylated peptidic chains were visualized by staining using an anti-puromycin antibody. The measured band intensity values were normalized to total protein values and shown as arbitrary units (a.u.).

### Human phospho-MAPK array

Kinase arrays were conducted as described in^59^. Briefly, 400 µg of protein from mock and HCV-infected Huh7 cells lysed at 72 h post infection or Huh7 cells treated for 1 h with DMSO or thapsigargin were analyzed using the Proteome Profiler Human Phospho-MAPK Array (R&D Systems) as recommended by the manufacturer. Signal was detected on a Vilber-Loumat imager and quantified using Image J software.

### Statistical analysis

Data from biological repeats is depicted as individual replicates with mean and standard deviation (SD). For experiments using multiple KO and Ctrl cell clones, data points and mean ± SD indicate variance among individual clones. Statistical analysis was performed using GraphPad Prism software, version 8.0.1. For comparison of two datasets, unpaired two-sided Student’s t-test was applied, using Welsh’s correction if datasets showed unequal variances. For comparison of three or more datasets with one or two variables, unpaired, two-sided one-way or two-way ANOVA was performed, respectively. To compare mRNA stabilities among datasets, statistical significance (ANOVA) over all time points between controls and stressed or KO conditions in indicated. For subsequent multiple comparisons, Dunnett’s posttest was applied for testing against a single control. For comparisons between multiple different groups of data, Holm-Sidak’s posttest was applied. P-values of less than 0.5 were considered statistically significant and are indicated accordingly (p < 0.05 = * or ^+^, p < 0.01 = ** or ^++^, p < 0.001 = *** or ^+++^, p < 0.0001 = **** or ^++++^).

---

**Figure S1.**
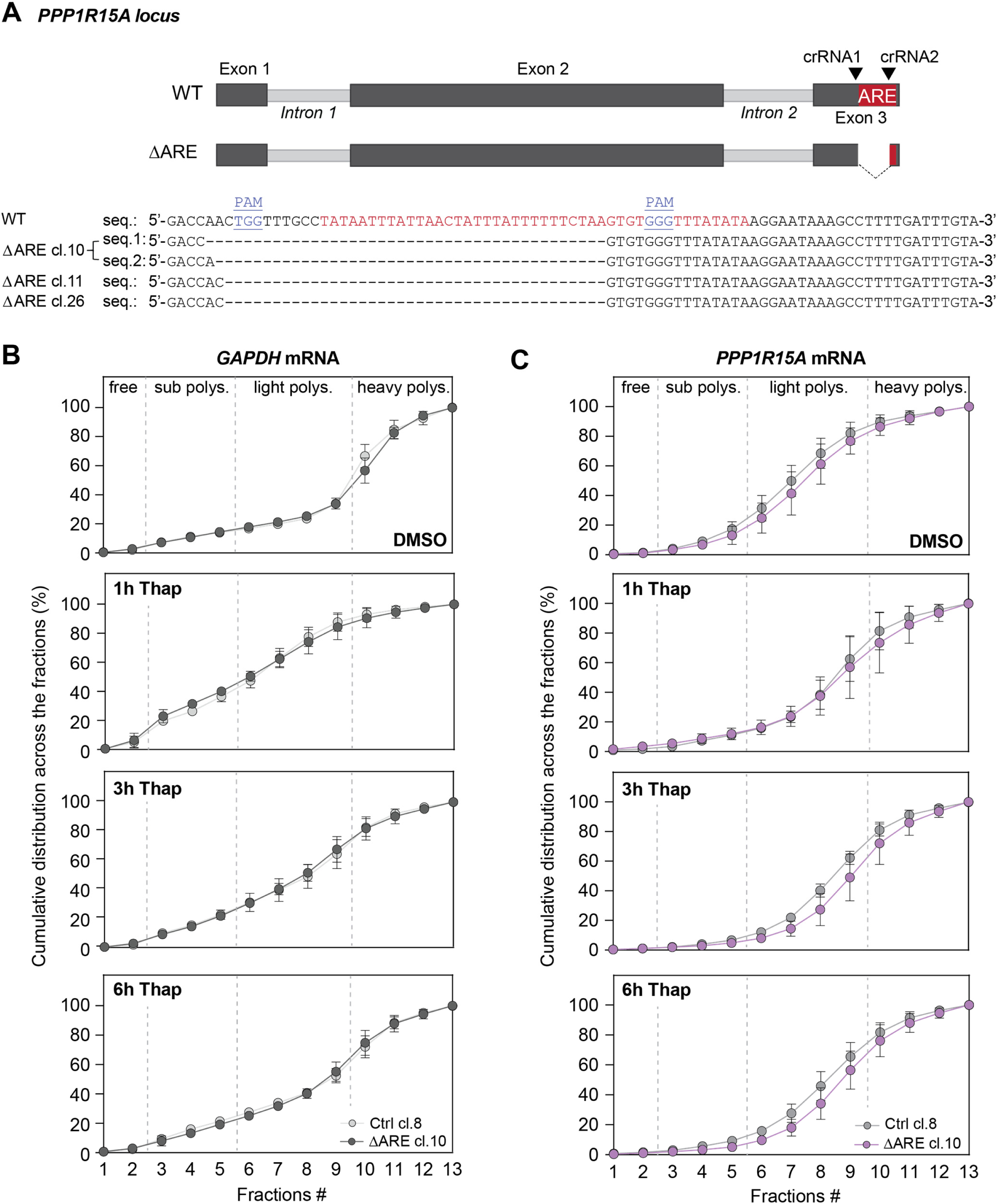
*PPP1R15A* ARE regulates mRNA engagement with polysomes. **Related to Figure 2. (A)** Generation of GADD34ΔARE cell clones. The top panel depicts PPP1R15A locus (WT). The AU-rich element (ARE) was deleted using two guide RNAs (crRNA1 and crRNA2) targeting *PPP1R15A* exon 3 (ΔARE). Three homozygous clones were selected based on the deletion of the genomic DNA sequence between the guide RNAs (PAM motifs are indicated in blue) and confirmed by sequencing. Sequences of individual alleles (up to two) for each clone are indicated. Nucleotide deletions are indicated by red dashes, insertions are marked in green. **(B and C)** Polysome profiles of Huh7 Ctrl cl.8 and Huh7 GADD34ΔARE cl.10 treated with thapsigargin for up to 6 h were recorded followed by fractionation of the sucrose density gradients. Shown are mean ± SD of two technical replicates from two distinct biological replicates (n=2). Total RNA was extracted from the polysome fractions, and the distribution of *PPP1R15A* and *GAPDH* mRNA across the polysome gradients was analyzed by RT-qPCR. The mean percentage ± SD of mRNA in each fraction is depicted as a cumulative distribution. The cumulative *GAPDH* mRNA distribution is shown in (B). Note that panel 6h Thap is the same as shown in Figure 2D. The cumulative *PPP1R15A* mRNA distribution is shown in (C).

**Figure S2.**
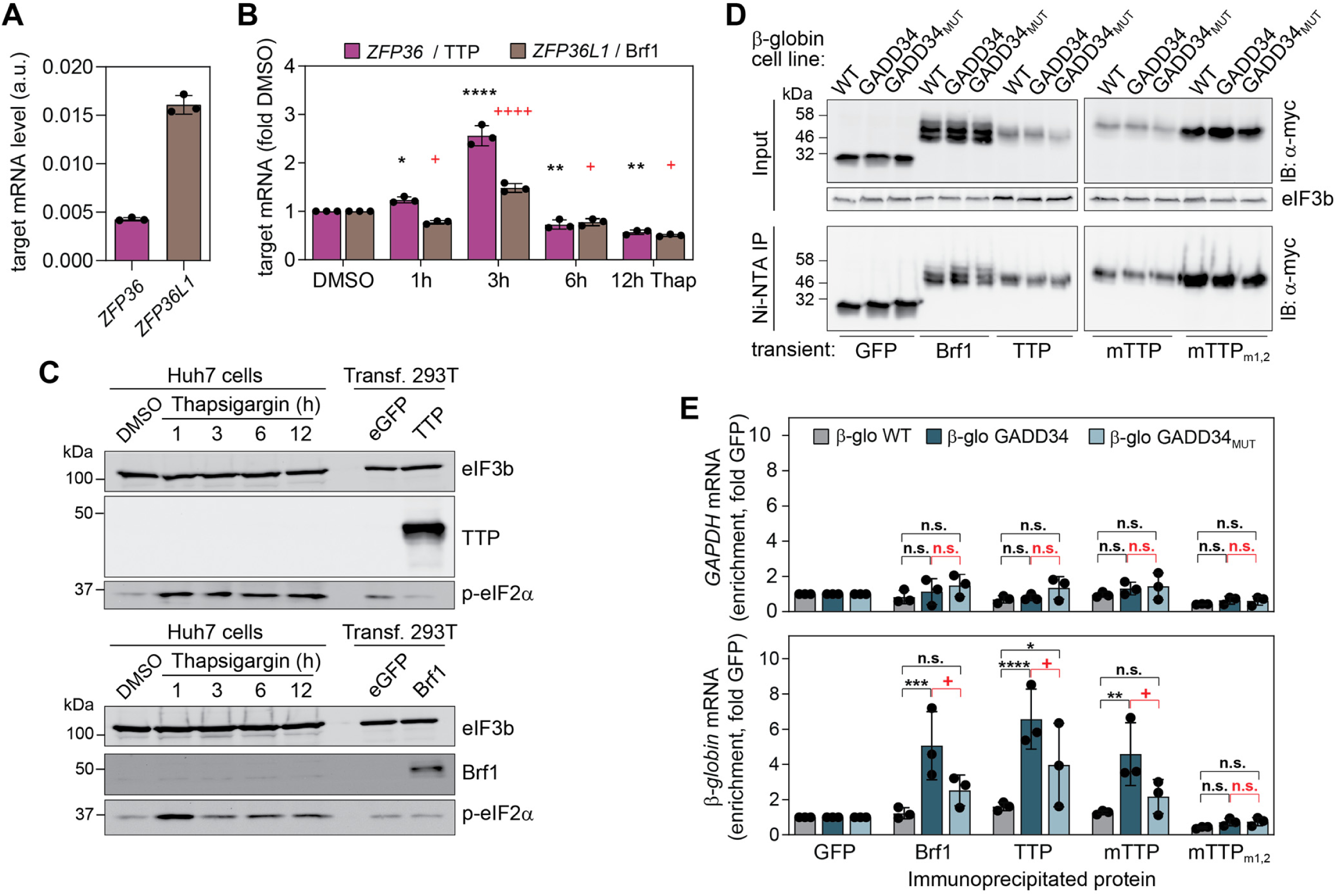
Expression of TTP and BRF1. **Related to Figure 4. (A and B)** Dynamic expression of *ZFP36* and *ZFP36L1* mRNAs. (A) Basal mRNA expression in Huh7 cells (mean ± SD, n=3). Values were normalized to *GAPDH* mRNA level values and depicted as arbitrary units (a.u.). **(B)** Relative mRNA expression in Huh7 cells treated with DMSO or thapsigargin for the indicated times (mean ± SD, n=3). Values were normalized to *GAPDH* mRNA level values and shown as fold DMSO. *, statistical significance compared to *ZFP36* mRNA levels in DMSO. +, statistical significance compared to *ZFP36L1* mRNA levels in DMSO. **(C)** Expression of endogenous TTP and Brf1 in Huh7 cells treated with DMSO or with Thapsigargin for up to 12 h. Shown is a representative Western blot for TTP expression (top panel) and Brf1 (bottom panel). Cell lysates of HEK293T cells overexpressing myc-his-tagged TTP and Brf1, respectively, were used as control. Levels of p-eIF2α served as control for ISR activation. eIF3b served as loading control. **(D and E)** The RNA-binding proteins TTP and Brf1 specifically bind to *PPP1R15A* mRNA ARE. Myc-his-tagged GFP, TTP, Brf1, TTP murine orthologue (mTTP) or RNA-binding mutant (mTTP_m1,2_) were immunoprecipitated using Ni-NTA beads from transiently expressing cells. RNA was subsequently isolated and analyzed by RT-qPCR (Ni-NTA-IP) (n=3). Upper panel: representative Western blot analysis. Shown are input cell extracts (10% of total, input) and immunoprecipitated proteins (50% of eluate, IP). Proteins were detected using an anti-myc antibody. eIF3b served as loading control. (E) Analysis of co-precipitated *GAPDH* (top panel) and *β-globin* (bottom panel) mRNAs, normalized to GFP. *, statistical significance compared to β-glo WT cells. **^+^**, statistical significance compared to β-glo GADD34 cells.

**Figure S3.**
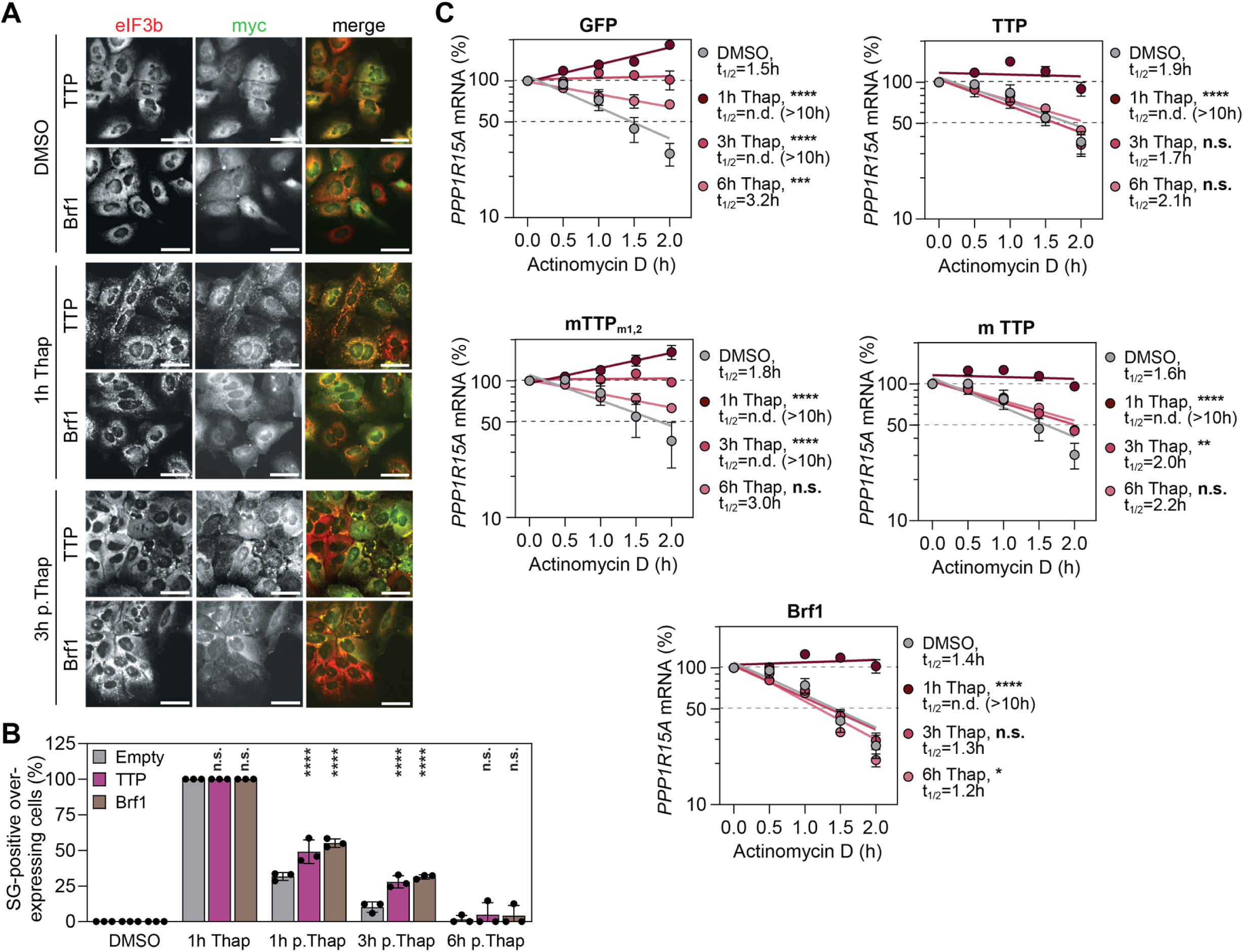
TTP and Brf1 overexpression interferes with *PPP1R15A* mRNA stabilization during stress adaptation and delay SG disassembly. **Related to Figure 4. (A and B)** Impact of ZFP36 proteins on SG disassembly kinetics. Huh7 cells transiently expressing myc-his tagged TTP and Brf1 were treated with thapsigargin for 1 h before drug removal and subsequent incubation in complete DMEM. Cells transduced with lentivirus carrying an empty viral vector (empty) were used as control. Cells were fixed after the 1-hour thapsigargin treatment (1h Thap) or at different time points during the stress adaptation phase, i.e. 1, 3 and 6 h post thapsigargin removal (p.Thap). (A) Immunofluorescence analysis of SG. TTP and Brf1 overexpressing cells were stained with the anti-myc-tag antibody (green). SG were stained with eIF3b (red). Shown are representative immunofluorescence images. Scale bar, 50 µm. (B) Corresponding analysis. Shown is the mean percentage of SG-positive cells ± SD (n=3). Statistical significance compared to control cells (transiently expressing the empty lentiviral vector) is indicated. (C) *PPP1R15A* mRNA decay in Huh7 cells transiently expressing myc-his tagged proteins and treated with DMSO or thapsigargin for the indicated times prior to actinomycin D chase. The original data depicted here is identical with Figure 4G but organized by cell type. Shown are mean ± SD (n=3). Statistical significance over all time points compared to DMSO is indicated.

**Figure S4.**
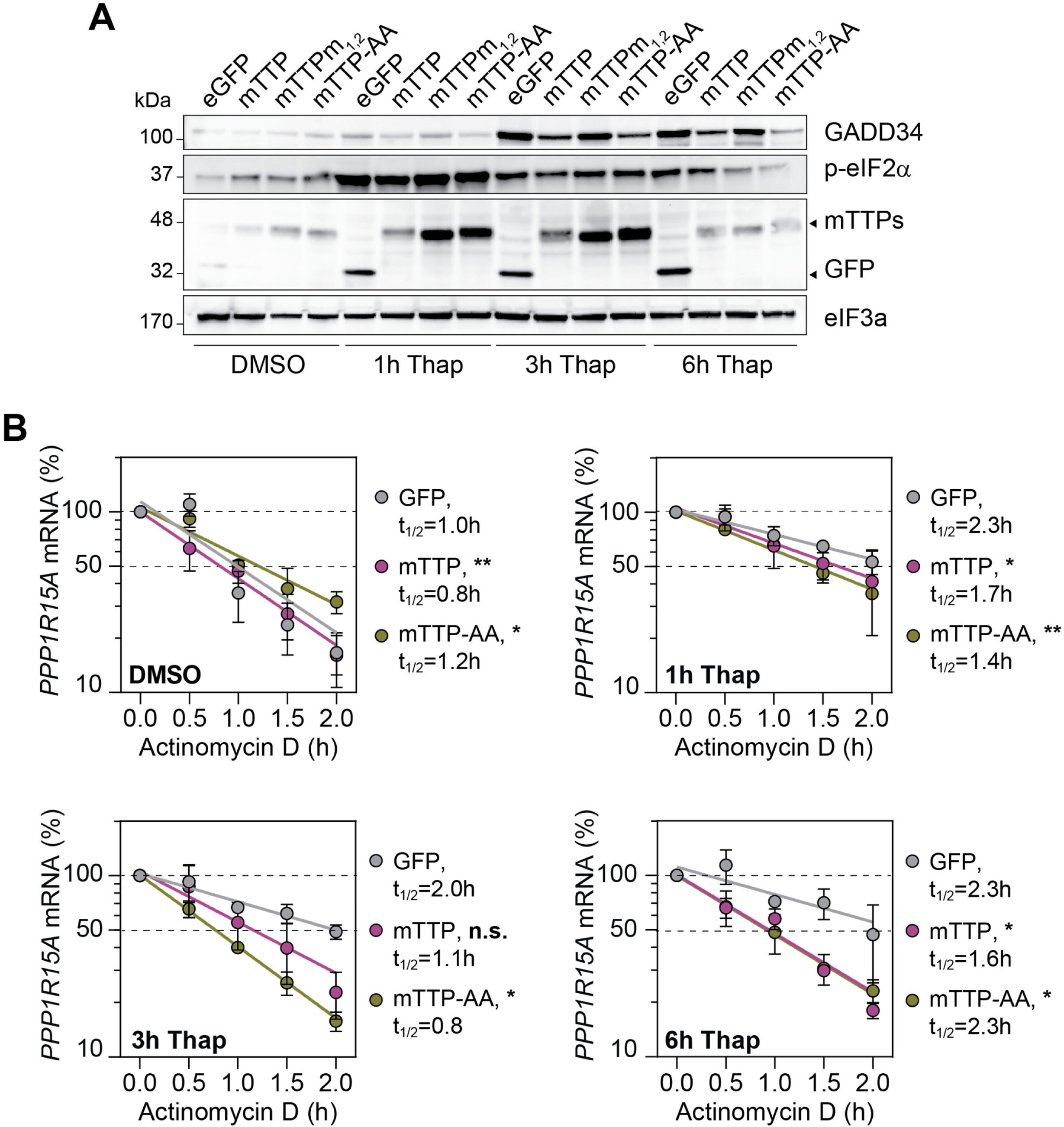
Phosphorylation-dependent inhibition of TTP contributes to the stabilization of *PPP1R15A* mRNA. **Related to Figure 4.** Myc-his-tagged GFP, mTTP, and the non-phosphorylable S52A/S178A mutant of murine TTP, mTTP-AA, were transiently expressed in Huh7 cells and subsequently treated with DMSO and thapsigargin for the indicated time. **(A)** Representative Western Blot analysis (n=3) showing myc-his-tagged protein expression levels, GADD34 and p-eIF2α. eIF3a served as loading control. **(B)** *PPP1R15A* mRNA decay in the corresponding cells. Shown are mean ± SD (n=3). Statistical significance over all time points compared to GFP is indicated.

**Figure S5.**
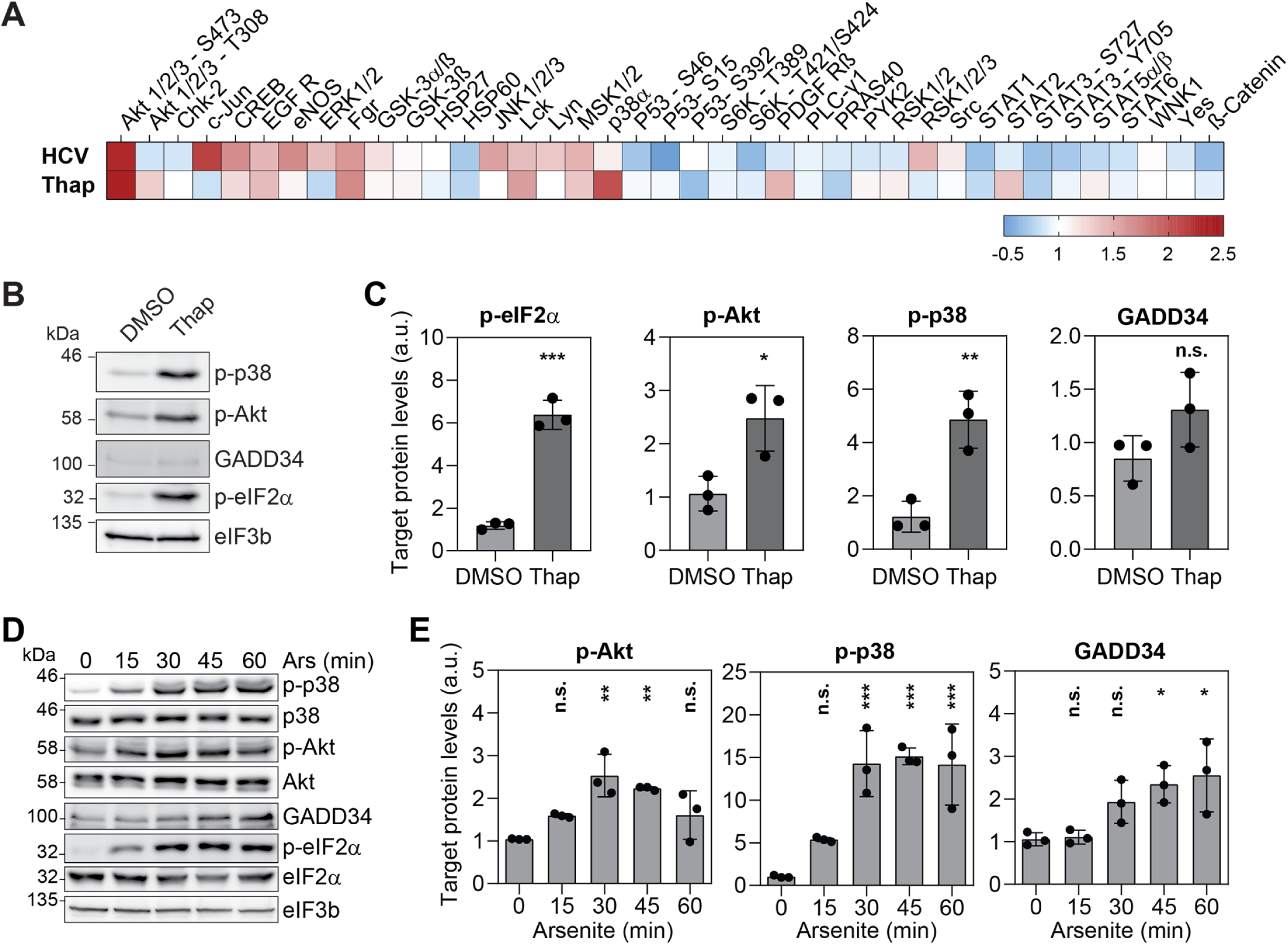
Transient activation of the p38-MAPK/MK2 and PI3K/Akt signaling pathways upon stress. **Related to Figure 5. (A)** The phosphorylation levels of several human MAPK kinases were measured using the human phospho-MAPK array in response to HCV infection in the presence of IFN-α or in response to thapsigargin treatment. Huh7 cells treated with IFN-α or with DMSO served as control, respectively. The heat map depicts changes in kinase phosphorylation levels compared to controls. Shown are means of independent biological replicates (n=3). Color gradient indicate fold changes compared to the corresponding controls. **(B and C)** Analysis of p38-MAPK/MK2 and PI3K/Akt signaling pathways in Huh7 cells treated with DMSO or thapsigargin for 1 h (n=3). Shown is a representative Western blot analysis (B) and corresponding quantifications of protein abundance (C). Measured band intensities were normalized to loading control (eIF3b) and depicted as arbitrary units (a.u.). Shown are means ± SD. Statistical significance compared to DMSO is indicated. **(D and E)** Shown is a representative Western blot analysis (D) and corresponding quantifications of protein abundance (C). Measured band intensities were normalized to loading control (eIF3b) and depicted as arbitrary units (a.u.). Shown are means ± SD. Statistical significance compared to T0 is indicated.

**Figure S6.**
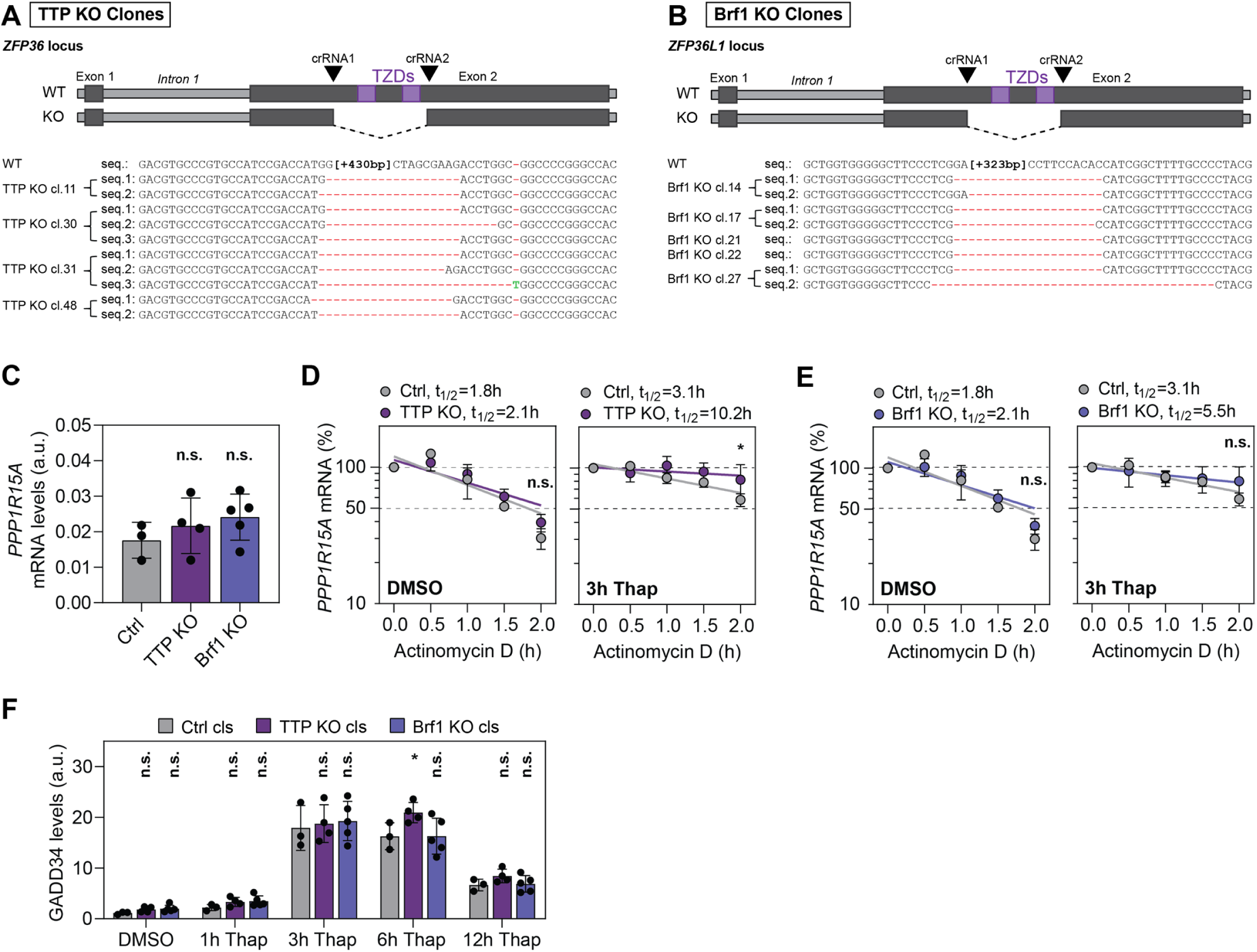
Characterization of TTP and Brf1 single KO cell clones. **Related to Figure 6. (A and B)** Huh7 KO cell clones were generated using the CRISPR/Cas9 two-guide RNAs approach (crRNA1 and crRNA2) to delete the region containing the TZF domains (TZDs) in exon 2 of the *ZFP36* and *ZFP36L1gene* loci. Homozygous KO clones were selected based on their genomic DNA sequence. Sequences of individual alleles (up to three) for each clone are indicated. Nucleotide deletions are indicated by red dashes. **(C)** Basal *PPP1R15A* mRNA expression in Huh7 TTP and Brf1 single KO cell clones. *PPP1R15A* mRNA levels were normalized to GAPDH mRNA levels and depicted as arbitrary units (a.u.). Statistical significance compared to Ctrl clones is indicated. **(D and E)** *PPP1R15A* mRNA decay measurements in single Huh7 KO clones. Three Ctrl, four TTP (D) and five Brf1 (E) cell clones were treated with DMSO or thapsigargin for 3 h (D and E) for 3 h prior to actinomycin D chase. Shown is the mean ± SD of all individual cell clones (n=1). Statistical significance over all time points compared to Ctrl clones is indicated. **(F)** Quantification of GADD34 expression levels in Ctrl, TTP KO and Brf1 KO single cell clones after exposure to DMSO or thapsigargin for the indicated times (n=2). Measured band intensities were normalized to loading control (eIF3b) and are depicted as arbitrary units (a.u.). Shown are representative means of cell clones ± SD. Statistical significance compared to Ctrl clones for each time point is indicated.

**Figure S7.**
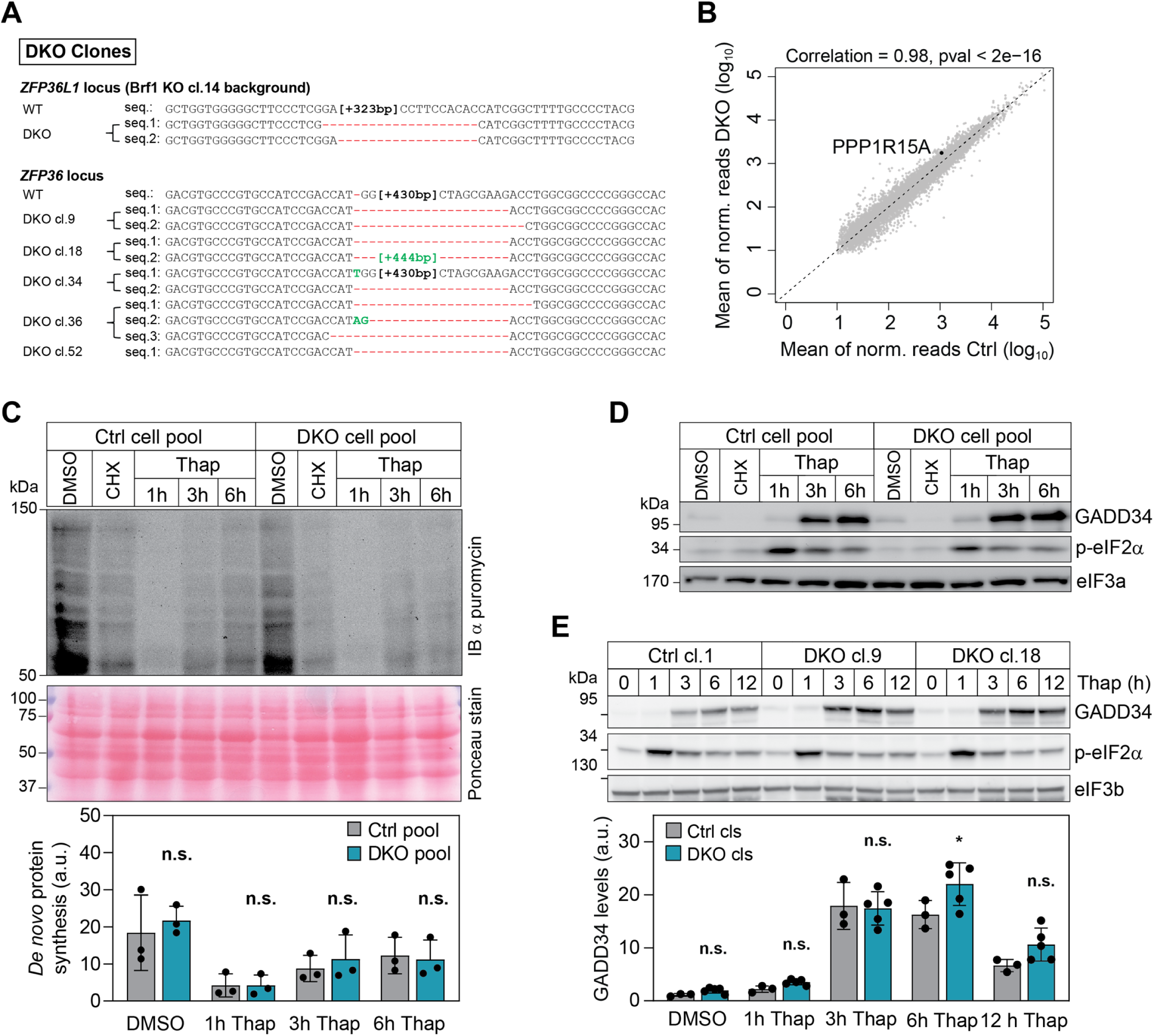
Characterization of Brf1 and TTP double KO (DKO) cell clones. **Related to Figure 6. (A)** Huh7 DKO cell clones were generated by deleting TTP TZD genomic region in Huh7 Brf1 KO cell clone 14. Homozygous KO clones were selected based on their genomic DNA sequence. Sequences of individual alleles (up to three) for each clone are indicated. Nucleotide deletions are marked by red dashes, insertions are marked in green. **(B)** Transcriptome-wide analysis of Huh7 Ctrl cell clone 1 and Huh7 DKO cell clone 18 (n=2). The scatter plot of the mean of normalized read counts (log 10) shows the correlation between Huh7 Ctrl cell clone 1 and Huh7 DKO cell clone 18. **(C)** Analysis of *de novo* protein synthesis in Huh7 Ctrl and DKO pooled clones treated with DMSO or thapsigargin for the indicated time (n=3). Shown is a representative Western blot analysis (top panel). Released puromycylated peptide chains were visualized by staining with an anti-puromycin antibody. The measured band intensity values were normalized to total protein values (ponceau stain, middle panel) and shown as arbitrary units (a.u.) (bottom panel). Shown is the mean ± SD (n=3). Statistical significance compared to Ctrl Pool for each time point is indicated. **(D)** Representative Western blot analysis of GADD34 and p-eIF2α expression levels in Huh7 Ctrl pooled clones and DKO pooled clones treated with DMSO or thapsigargin for the indicated time (n=4) (related to Figure 6C and 6D). eIF3a was used as loading control. **(E)** Representative Western blot analysis of GADD34 and p-eIF2α expression levels in Ctrl clone 1 and DKO clones 9 and 18 upon thapsigargin treatment (n=2). eIF3b was used as loading control. Related to Figure 7D.

**Figure S8.**
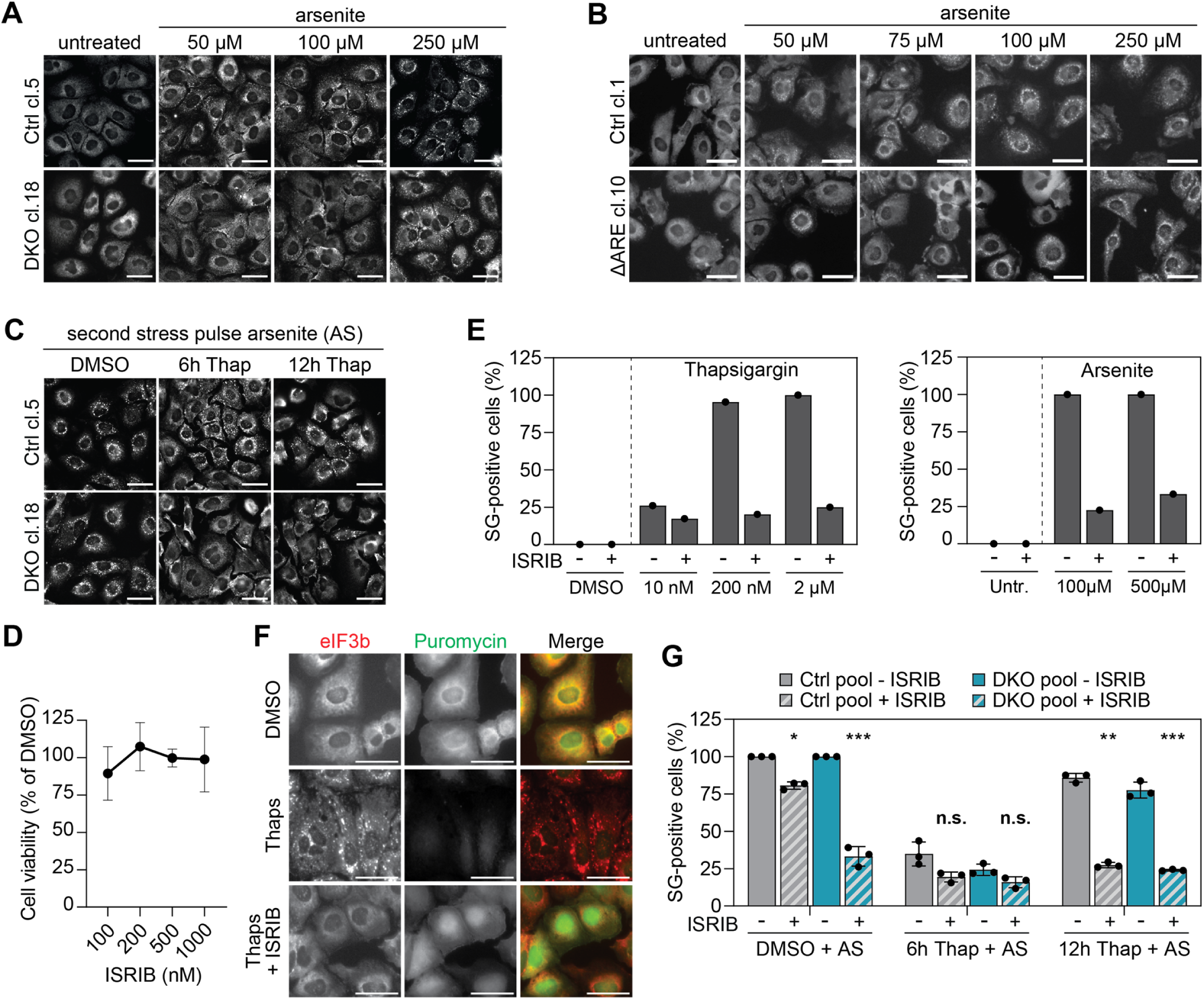
ZFP36 proteins contribute to set the threshold for stress responsiveness and the dynamics of stress adaptation. **Related to Figure 7. (A and B)** Analysis of cell responsiveness to increasing doses of arsenite (related to Figures 7A and 7B). The ability of cells to form SGs was assessed by immunofluorescence analysis. Cells were fixed and stained for eIF3b as SG marker. (n=3). Shown are representative immunofluorescence microscopy images showing SG formation in Huh7 Ctrl cl.5 and Huh7 DKO cl.18 (A) and in Huh7 Ctrl cl.1 and Huh7 GADD34ΔARE cl.10 (B). Scale bar, 50 μm. **(C)** Huh7 Ctrl and DKO cell clones were treated with DMSO or thapsigargin for 6 or 12 h before a second stress pulse with 500 µM arsenite for 1 h (acute stress, AS) (related to Figures 7E). Cells were fixed and stained for eIF3b as SG marker. Shown are representative immunofluorescence images for Ctrl cl.5 and DKO cl.18. Scale bar, 50 μm. **(D to F)** Characterization of ISRIB inhibition in Huh7 cells (n=1). (D) Cell viability after 1 h treatment with the indicated concentration of ISRIB determined by WST-1 assay and represented as percentage of DMSO-treated cells. (E) Percentage of SG-positive Huh7 cells upon treatment with increasing concentration of thapsigargin (left panel) or sodium arsenite (right panel) in the absence and presence of 1000 nM ISRIB (n=1). (F) Analysis of *de novo* protein synthesis by puromycylation assay in Huh7 treated with DMSO, 2 µM thapsigargin for 1h (Thap), or co-treated with 2 µM thapsigargin and 1000 nM ISRIB for 1h (Thap + ISRIB). Shown are representative immunofluorescence images. Cells were fixed and stained for eIF3b as SG marker. Released puromycylated peptide chains were visualized by staining with an anti-puromycin antibody. Scale bar, 50 µm. **(G)** ISRIB inhibits the ISR and SG formation in response to acute stress and during the adaptation phase. Analysis of SG formation in Huh7 Ctrl and DKO cell clone pools pre-treated with DMSO and thapsigargin for 6 and 12 hours prior a second acute stress pulse with 500 µM arsenite, in the presence and absence of 1000 nM ISRIB. Shown is the mean percentage of SG-positive cells ± SD (n=3). Statistical significance compared to absence of ISRIB (-ISRIB) is indicated.

**Figure S9.**
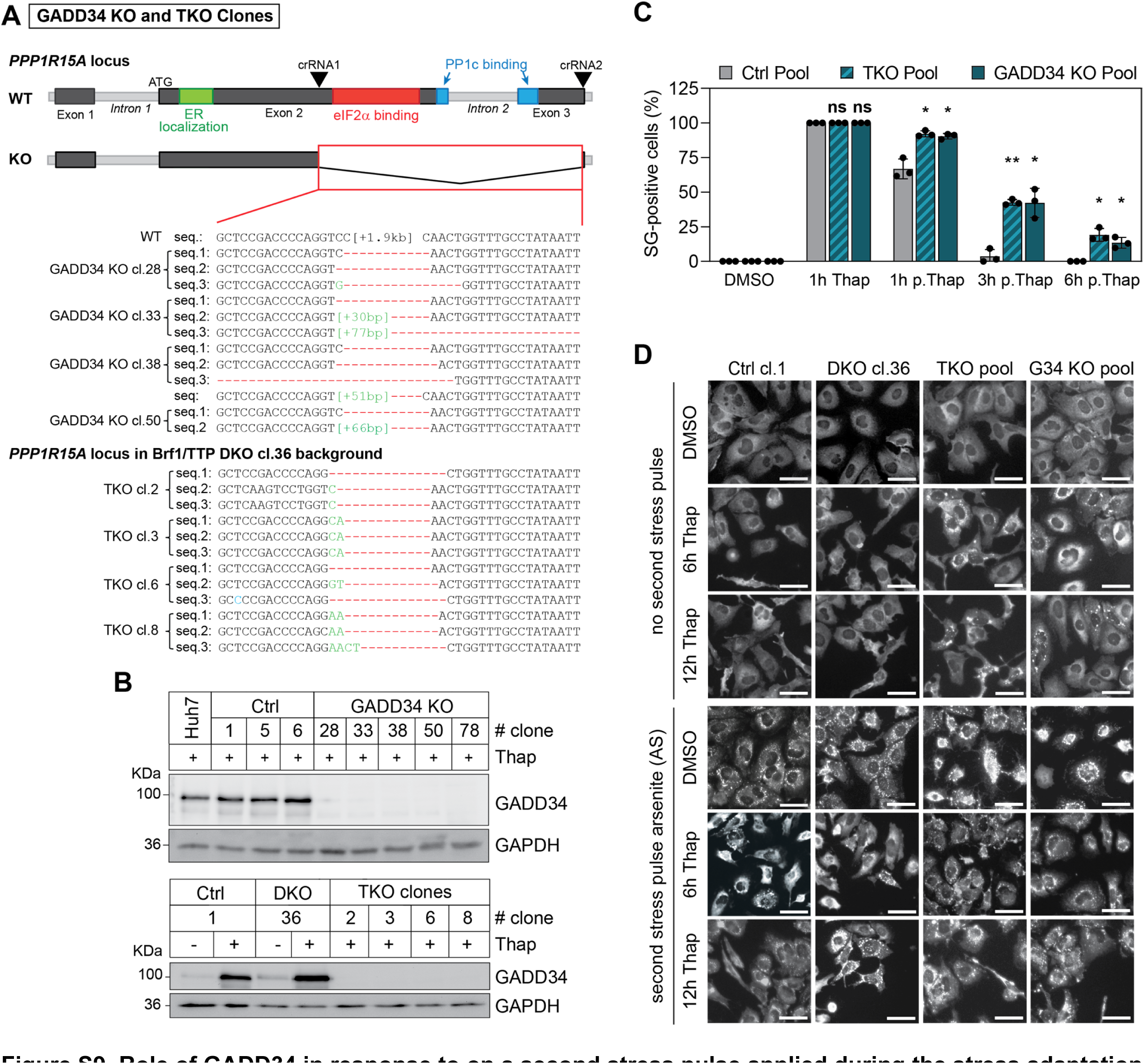
Role of GADD34 in response to on a second stress pulse applied during the stress adaptation phase. **Related to Figure 7. (A and B)** Characterization of Huh7 GADD34 KO and Brf1/TTP/GADD34 (TKO) cell clones. (A) Huh7 GADD34 KO were generated by deleting a sequence in *PPP1R15A* genomic locus using a CRISPR/Cas9 two-guide RNAs approach (crRNA1 and crRNA2) to delete the region containing eIF2α and PP1c binding domains in exons 2 and 3. Huh7 TKO cell clones were generated by deleting the same *PPP1R15A* sequence in the TTP/Brf1 DKO cell clone 36 background. Homozygous KO clones were selected based on their genomic DNA sequence. Sequences of individual alleles (up to three) for each clone are indicated. Nucleotide deletions are marked by red dashes, insertions are marked in green. (B) Analysis of GADD34 expression in control (Ctrl), GADD34 KO cell clones (top panel), and TKO clones (bottom panel) treated with thapsigargin for 3 h. GAPDH served as a loading control. **(C)** Analysis of SG disassembly in Ctrl, TKO and GADD34 KO cell clone pools during stress adaptation. Cells were fixed after the 1-hour thapsigargin treatment (1h Thap) or at different time points during the stress adaptation phase, i.e. 1, 3 and 6 h post thapsigargin removal (p.Thap). Shown is the mean percentage of SG-positive cells ± SD (n=3). Statistical significance compared to Ctrl cell clones at the respective time point is indicated. **(D)** Impact of GADD34 during the stress adaptation phase in response to on a second stress pulse. Huh7 Brf1/TTP/GADD34 triple knockout (TKO) and GADD34 knockout (G34 KO) cell pools were compared to Huh7 Ctrl cl. 1 and Huh7 DKO cl.36. Cells were treated with DMSO or thapsigargin for 6 and 12 h and subsequently subjected to a second stress pulse of high dose arsenite for 45 min (acute stress, AS). The ability of cells to form SGs was assessed by immunofluorescence analysis (related to Figure 7F). Cells were fixed and stained for eIF3b as SG marker (n=3). Shown are representative immunofluorescence microscopy images of cells prior to acute stress (top panel) and after acute stress (bottom panel). Scale bar, 50 μm.

